# Parallel Evolution of Bacteroidota as Long-Term Endosymbionts of Insects

**DOI:** 10.1101/2025.10.31.685726

**Authors:** Jinyeong Choi, Cong Liu, Pradeep Palanichamy, Yumiko Masukagami, Hirotaka Tanaka, Takumasa Kondo, Matthew E. Gruwell, Filip Husnik

**Affiliations:** Evolution, Cell Biology, and Symbiosis Unit, Okinawa Institute of Science and Technology Graduate University, Okinawa 904-0495, Japan; Evolutionary Genomics Unit, Okinawa Institute of Science and Technology Graduate University, Okinawa 904-0495, Japan; Faculty of Agriculture, Ehime University, Tarumi 3-5-7, Matsuyama, Ehime 790-8566, Japan; The Kyushu University Museum, Hakozaki 6-10-1, Higashi-ku, Fukuoka, 812-8581 Japan; Corporación Colombiana de Investigación Agropecuaria-Agrosavia, Centro de Investigación Palmira, Valle del Cauca, Colombia; Penn State Erie, Behrend College, School of Science, Erie, PA, USA

## Abstract

Symbiotic relationships transform diverse aspects of both symbiont and host biology. The most visible changes include a massive reduction of the endosymbiont genome and the development of novel host organs, cells, and compartments specialized for harboring the symbionts. However, many insect symbiosis studies have previously focused either on Proteobacteria or only on a particular symbiosis stage, limiting our broad understanding of how and why symbiont diversity arises from specific microbial clades, how the symbiont genomes erode over evolutionary time, and what the consequences of symbiosis are for diverse host-symbiont pairs. Thanks to the repeated gains and losses of nutritional symbionts, scale insects provide an ideal evolutionary playground for tracking the parallel transitions of insect symbionts that originated as independent evolutionary replicates from the same bacterial phylum. Using extensive genome sampling across the scale insect phylogeny, we recapitulated the path of Bacteroidota transitioning from recently established host-associated bacteria to highly specialized insect endosymbionts with minimal gene sets. Their genomes exhibit strikingly parallel patterns of gene loss and pseudogenization across all gene categories, with some stochastic differences in essential genes for genetic processing and amino acid biosynthesis. In addition to the genetic transition, we show with imaging methods that the symbiotic cells and organs exhibit a trend from more dispersed bacteriocytes to highly compact (and likely more specialized) bacteriomes, which are closely localized to the host’s digestive system. Our results outline at both genomic and cellular levels a recurrent path by which insect symbioses independently arise, are temporarily maintained for up to several hundred million years, and then are replaced by new symbionts that repeat the process. The gradual nature of the process implies that the outcomes of symbiosis initially depend on how ‘professional vs. naïve’ the host and symbiont are. Over evolutionary time, compatible host-symbiont lineages emerge from these interactions and become preferred.

## Introduction

The evolutionary fate of microbial symbionts is predetermined, and in the case of maternally transmitted symbionts, it is almost irreversible due to population genetics processes^1^. With ongoing gene loss due to the relaxed selection and random genetic drift accelerated by intrahost localization and bottlenecking during transmission^2^, most symbiont genes are eliminated over time. Genes that are erased last are involved in DNA replication, transcription, and translation, as well as in a few pathways that confer benefits to the host^3,4^. This further pseudogenization of the important symbiont genes is caused by accidental early stop codons arising from nonsense mutations^5^. To alleviate this disruption in protein-coding genes, the old symbiont genomes have occasionally undergone codon reassignments, resulting in an alternative genetic code that translates the UGA codon to tryptophan instead of its original role as a translational stop signal^7^. In many cases, the symbiont functions have been shown to be replaced or complemented by host genes of both foreign and native origin or by additional co-obligate symbionts^8,9,10,11^. However, despite these processes slowing down symbiont degradation, most long-term endosymbionts face extinction, replacement by new microbial partners, or, in some cases, organelle-like integration^12,13^.

Bacteroidota play critical roles in nutrient cycling in both aquatic and terrestrial ecosystems^14^. Many Bacteroidota are also crucial symbionts of animals (including humans), plants, and many other eukaryotes. Their diverse ecological niches and multifunctional roles are reflected in their high genomic diversity and plasticity^14^. In insects, Bacteroidota are well-known as beneficial symbionts establishing long-term evolutionary relationships with diverse insect hosts^4,15,16,17,18^. These relationships have been proposed to expand the ecological niches of the host insect groups and promote their diversification^19^. A textbook example is *Candidatus* Karelsulcia, which has been maintained by auchenorrhynchan insects for about 300 million years, and its strains exhibit a substantial genome reduction (142–285 kb)^15,20,21^. In terms of genome size, *Karelsulcia* is followed by *Candidatus* Shikimatogenerans (172–200 kb) and *Candidatus* Bostrichicola (323– 346 kb) in bostrichid beetles^4^, *Blattabacterium* (511–645 kb) in cockroaches and termites^22,23^, and *Candidatus* Skilesia (1.32 Mb) in aphids^17^. However, the sampling of Bacteroidota symbionts in insects is limited to only one or two lineages per insect group, and only when combined, they represent multiple stages of genome reduction, highlighting the importance of broader sampling specific to a single insect clade.

Scale insects house long-term obligate symbionts that provide them with essential amino acids and vitamins lacking in their plant sap diets^26,27^. In contrast to other insect clades with Bacteroidota symbionts, scale insects have a dynamic evolutionary history with various lineages of Bacteroidota symbionts^26,28,29^. Despite the diversity of Bacteroidota lineages recognized as associated with scale insects, genome sequencing and microscopy have been limited to only two genera of Bacteroidota symbionts, *Candidatus* Walczuchella (270–309 kb)^30,31^ and *Candidatus* Uzinuria (263 kb)^32,33^. Here, we analyzed 21 genomes of 10 different Bacteroidota lineages from diverse scale insect clades. Our paired host-symbiont phylogenomic analyses robustly show that scale insects have acquired Bacteroidota symbionts multiple times independently. If there ever was an ‘ancient’ Bacteroidetes symbiont in the ancestor of all or most scale insects, it has been replaced over evolutionary history. We reveal significant variation in the symbiont genome sizes and gene contents, suggesting an ongoing process of pseudogenization and gene loss that depends on the age of the symbiont. Additionally, we show with microscopy that, depending on the symbiont genome reduction level, the localization of symbiont-harboring cells and organs tends to shift from outer body regions and fat-body cells to the close proximity of the insect gut. These results contribute to the understanding of the evolutionary dynamics of Bacteroidota, showcasing how they frequently transition into highly integrated symbionts of insects, and suggest how compatible host-symbiont lineages evolve over hundreds of millions of years.

## Materials & methods

### DNA extraction and metagenome sequencing

Metagenomes of 18 species were newly generated in this study. Insect samples were preserved in 99% ethanol at –20°C. Total genomic DNA was extracted with the MasterPure Complete DNA Purification Kit (Epicenter) and DNeasy Blood and Tissue Kit (Qiagen) following the manufacturer’s protocols. PCR-free sequencing libraries were prepared with the NEBNext Ultra II DNA library preparation kit. In addition to newly generated data, the Sequence Read Archive (SRA) data from 23 species were reused in this study. In total, we analyzed metagenomes from 41 species in 16 families of scale insects (Supplementary Table 1). Paired-end sequencing was performed on the Illumina NovaSeq 6000 sequencer at the Okinawa Institute of Science and Technology Graduate University and MiSeq at SEQme and Penn State University. We confirmed the quality of raw Illumina reads using FastQC v0.11^34^ and filtered out low-quality reads with fastp v.0.20^35^. The taxonomic affiliation and relative abundance of different organisms were profiled using phyloFlash v3.4^36^ based on 16S and 18S rRNA gene sequences from the metagenomes. All commands used for the bioinformatics analyses are available from the following GithHub repository: https://github.com/ECBSU/ECBSU_manuscripts_code/tree/main/Scale_Insects_Bacteroidota.

**Table 1.** Features of select animal symbionts using the genetic code with UGA: stop −> tryptophan.

| Class | Flavobacteriia | α-proteobacteria | β-proteobacteria | | γ-proteobacteria | |
| --- | --- | --- | --- | --- | --- | --- |
| Symbiont | Bacteroidota<br>RAIC | *Hodgkinia* | *Nasuia* | *Zinderia* | *Stammera* | *Providencia* |
| *prfA* | - | + | + | + | + | + |
| *prfB* | - | - | - | - | - | +($\psi$)* |
| *trnW* anticodon | UCA | UCA | UCA | CCA | CCA | CCA |
| GC% | 35.8 | 58.4 | 17.1 | 13.5 | 15.4 | 22.0 |
| CDSs | 185 | 169 | 137 | 206 | 255 | 610 |
| In-frame UGA (%) | 101 (55%) | 119 (70%) | 69 (50%) | 121 (59%) | 159 (62%) | 125 (21%) |

\*Including a UGA stop codon in the *prfB* pseudogenes.

### Phylogenetic analyses of symbionts

Phylogenetic trees of symbionts were reconstructed using the rRNA gene sequences and single-copy orthologous genes. The rRNA gene phylogeny was primarily used to identify symbionts from metagenomes. The SSU (16S/18S) rRNA and LSU (23S/28S) rRNA gene sequences of bacterial and fungal symbionts were extracted from metagenome assemblies with Barrnap v3 (https://github.com/tseemann/barrnap). Additionally, we also used published genome assemblies and rRNA sequences of related symbionts and free-living bacteria from NCBI (accession numbers shown on phylogenetic trees). The SSU rRNA and LSU rRNA gene sequences were concatenated after individual alignment with MAFFT v7^37^ and trimming with trimAL v.1.4^38^ (-automated1). We used two different approaches for the phylogenomic analysis of Bacteroidota symbionts. First, we predicted orthologous genes using OrthoFinder v.2.5.4^39^ and extracted only 65 genes that were present as single-copy in all the Bacteroidota genomes. These protein sequences were aligned with MAFFT v7 (--auto), trimmed with trimAL v1.4 (-automated1), and then concatenated. Second, we sampled single-copy orthologous genes from genome assemblies of Bacteroidota with BUSCO v.5.1.3^40^ (with the bacteroidetes_odb10 database). The protein sequences of 118 orthologous genes were aligned with MAFFT v7 and concatenated with Seqkit^41^ after trimming each gene sequence with trimAL v1.4 (-automated1). Maximum likelihood analyses for all the sequence matrices above were performed with IQ-TREE v.1.6.9^42^. The substitution models were chosen with ModelFinder^43^ implemented in IQ-TREE (COMMAND). Node support was assessed with 1,000 ultrafast bootstrap pseudo-replicates. The phylogenetic trees were visualized using Figtree v1.4.4^44^. The following Candidatus status names have been assigned to Bacteroidota lineages that currently lack a scientific description. First, *Candidatus* Brownia xenocola is for the symbionts of *Eumyrmococcus scorpioides*. The species name *xenocola* refers to the host family, Xenococcidae. This bacterium belongs to the same genus as *Brownia rhizoecola*, a symbiont of Rhizoecidae, due to their close phylogenetic relationship. Second, *Candidatus* Conchaspibacter symbioticus is for the symbionts of *Conchaspis* sp. Third, *Candidatus* Orthebacter symbioticus is for the symbionts of *Orthezia urticae*. Fourth, *Candidatus* Insignorthebacter symbioticus for the symbionts of *Insignorthezia insignis*. Fifth, *Candidatus* Acanthobacter symbioticus is for the symbionts of *Acanthococcus lagerstroemiae*. Sixth, *Candidatus* Paraacanthobacter symbioticus is for the symbionts of *Acanthococcus* sp. The genus names are derived from the associated hosts, with “-bacter” indicating their bacterial affiliation, while the “symbioticus” species names reflect their symbiotic lifestyle within the hosts.

### Symbiont genome assembly and annotation

To obtain the draft genomes of Bacteroidota symbionts and one *Sodalis*-like co-symbiont, the raw Illumina reads were assembled using SPAdes v3.15^45^ with specific k-mer sets and options (Supplementary Table 2). In some cases, we used the merged reads generated by PEAR v.0.9.11^46^ and the untrusted-contigs option with available reference genomes. The assembled sequences were taxonomically assigned using megablast search (NCBI-BLAST v.2.11.0) against the NCBI nucleotide database with the parameters “-outfmt ‘6 qseqid staxids bitscore std sscinames sskingdoms stitle’-culling_limit 5-evalue 1e-25”. The taxon-annotated-GC-coverage plots were generated for each metagenome with Blobtools v1.1.1^47^. Based on these plots, the genome assemblies of Bateroidota symbionts were extracted from the metagenomes. Due to the high number of contigs of the Bacteroidota ORUR genome, we used MetaBAT2 v.2.12.1^48^ to bin the genome and assessed its completeness with the DASTool v.1.1.6^49^. Genome assemblies with 2 or 3 contigs (Bacteroidota RAIC, RASP, RHAM, and RIMU) were closed manually using the alignment with other assemblies and assembly graphs as a guide. The joint regions of the original contigs were confirmed by read mapping and visualization with Tablet v.1.21.02.08^50^. Because the Illumina-only assemblies of ACLA and COCA metagenomes resulted in fragmented symbiont genomes, we also assembled their PacBio reads using Flye v.2.9^51^ with the --meta option. Assembly graphs of all the Bacteroidota genomes were visually assessed with Bandage v.0.9^52^. All the genome assemblies of Bacteroidota were polished using Pilon v.1.24^53^. PacBio-only assemblies of Bacteroidota ACLA and COCA were polished with Illumina short reads available from NCBI. Genome annotations were carried out with Prokka v.1.14.6^54^. Some gene predictions were confirmed with Bakta v.1.8.1^55^ and GhostKOALA v.2.0^56^. Bacteroidota symbiont RAIC was found to be using an alternative genetic code, so we used the genetic code 4 (UGA > Trp) for gene prediction instead of the code 11 used for all the other symbiont genomes. Hypothetical proteins were re-annotated through BLASTp searches against the NCBI RefSeq (NR) database^57^. The transfer RNA genes (tRNAs) were predicted with tRNAScan-SE v2.0^58^. Missing intact CDSs in the gene prediction were recovered with BLASTx searches of intergenic regions, implemented in Pseudofinder v1.1.0^59^. The coding sequences (CDSs) were classified into clusters of orthologous groups (COGs) using eggNOG-mapper v2^60^ with default parameters. Orthologs of Bacteroidota genomes were inferred with OrthoFinder v.2.5.4^39^. Genome feature maps were created with DNAPlotter^61^. A synteny plot of Bacteroidota genomes was generated with Processing3 (https://processing.org) based on tBLASTx genome alignments and Prokka annotations.

### Pseudogene and selection pressure predictions

The putative pseudogenes of Bacteroidota symbionts were predicted with PseudoFinder v1.1.0 with DIAMOND v2.0.4.142^62^ searches against NR. We further estimated the selective pressure on the symbiont genes mostly involved in genetic information processing and AA biosynthesis. The nonsynonymous mutation rate to the synonymous mutation rate (*dN/dS*) of each orthologous gene was calculated using the yn00 program based on pairwise comparisons in PAML v.4.9^63^. The free-living bacterium, *Flavobacterium johnsoniae* (GCF_034479105), with non-reduced genomes (> 6 Mbp), was used as a standard for the sequence comparison. The orthologous gene pairs of each Bacteroidota symbiont and the free-living bacterium were identified using OrthoFinder. For genes with paralogs, we selected only one sequence of each bacterium based on its sequence similarity. The amino acid sequences of orthologous genes were aligned with MAFFT v7 (--auto). The codon-based DNA alignment was obtained with PAL2NAL v.14^64^ based on the aligned amino acid sequences and unaligned nucleotide sequences. We considered the genes with a *dN*/*dS* ratio ranging from 0.1 to 0.95 to be evolving under relaxed purifying selection^65^.

### Host genome assemblies and phylogenomic analyses

A custom pipeline was built to extract the host insect genomes from 39 metagenome assemblies. First, we removed contigs shorter than 400 bp and mapped the raw reads onto the filtered metagenomic assembly with minimap2^66^. Then we computed the sequencing depth, GC content, and coding density for each contig with SprayNPray VERSION ^67^. We also searched the contigs against the non-redundant (nr) database with DIAMOND VERSION ^68^ and assigned a putative phylum to each contig with MEGAN VERSION ^69^ if available. Second, we trained a decision tree classifier using scikit-learn, with the sequencing depth, GC content, and coding density as training features and phylum as the target value. This classifier inferred phyla for contigs that were not classified by MEGAN. Finally, we extracted contigs assigned to Arthropoda and manually removed contigs with distinct depth and GC content from the main cluster. The extracted insect genome bin was evaluated by QUAST^70^ and BUSCO (with the hemiptera_odb10 database). A total of 57 species were used for phylogenomic analysis, including 18 published genomes retrieved from NCBI. A matrix of concatenated protein sequences was assembled from 359 single-copy genes identified by BUSCO VERSION following the methods as described above for the symbiont phylogenetic analysis. A maximum likelihood (ML) tree was produced with IQ-TREE under the best-fitting model (JTT+F+R6) selected automatically, and 1,000 ultrafast bootstrap replicates were performed to assess the node support. Divergence times were estimated using MCMCTree in PAML v.4.9^63^ with the independent rate model. The following minimum-age calibrations were used based on the oldest fossils to respective nodes of clades: (i) Ortheziidae (135 Ma)^71^ and (ii) Pseudococcidae (135 Ma)^72^. The root age was constrained to <300 Ma according to the fossil information and previous study^73,74^. We performed two independent runs (nsample = 10,000, sampfreq = 1,000, burnin = 1,000,000) to confirm the convergence of MCMC chains with Tracer v.1.7.2^75^. The mean and 95% confidence intervals were assessed. All the phylogenetic trees were visualized with FigTree.

### Microscopy

Localization of Bacteroidota symbionts in host insects was examined using an integrated approach of fluorescence in situ hybridization (FISH) and micro-computed tomography (µCT). We used three host species (ACLA, RASP, and RAIC), harboring Bacteroidota symbionts at early, intermediate, and late stages of symbiosis based on their genome status (Supplementary Table 1). For FISH, the distribution of symbionts was confirmed using a 16S rRNA probe CFB319 (Cy3-TGGTCCGTGTCTCAG-TAC), targeting Bacteroidota^15^. Whole insect tissues were fixed for 1 day in 4% paraformaldehyde in phosphate-buffered saline (PBS-1X) and Carnoy’s solution. After washing with 80% ethanol, tissues were bleached for 2–4 weeks in 6% hydrogen peroxide solution, which was replaced with fresh solution several times. After washing with absolute ethanol, the bleached tissues were embedded in Technovit 8100 resin (Kuzler, Wehereim, Germany) and sectioned into 3–5 μm sections using a rotary microtome (HM 340E, Epredia). The sections were incubated overnight at room temperature in hybridization buffer containing DAPI (1µg/µl) and probes (100 nM). The samples were mounted with ProLong Diamond Antifade Mountant (Thermo Fisher Scientific) after washing with PBS. Tissues were then examined under the Nikon ECLIPSE Ts2-FL (Nikon, Tokyo, Japan) inverted microscope. The three-dimensional distribution of symbionts was further confirmed using µCT. The samples were stained with iodine for 1–3 days and scanned using the Zeiss XRadia Versa410 X-ray microscope, operated with the Zeiss Scout-and-Scan Control System software (v.14.0). Scanning parameters included a beam strength of 40 kV and 3 W power, with exposure time of 13–19 s under 4x objective lens, resulting in voxel size of 1.09– 1.49 μm. Post processing was performed using Amira 2019.2 software (Visage Imaging GmbH, Berlin, Germany).

## Results

### Well-supported scale insect phylogeny with divergence time estimates

Draft genomes of 39 species of scale insects were newly assembled in this study (genome quality statistics in Supplementary Table 3). The average size and GC content of the assemblies were 315 Mbp and 33%, respectively. The average scaffold N50 of assemblies was 28 Kbp, and 24 genomes had N50 above 10 Kbp. The BUSCO completeness of 57 genomes, including 18 NCBI references, averaged 75%, with 39 genomes exhibiting completeness above 80%. Maximum likelihood analysis with IQ-TREE was well-resolved for most clades (Fig. 1; Supplementary Fig. 1). The Coccomorpha clade was estimated to have originated approximately 300 million years ago (Mya) (Fig. 1; Supplementary Fig. 2). Among hosts with long-term Bacteroidota symbionts, Monophlebidae, Rhizoecidae, and Diaspidiae species formed distinct clades. First, the Monophlebidae clade was estimated to have appeared approximately 140 (Mya) in the Cretaceous or possibly the Jurassic. Second, the Rhizoecidae clade was sister to the Pseudococcidae and was estimated to have originated 100 Mya in the Cretaceous. Third, the Diaspididae clade was estimated to have originated 50 Mya in the Paleogene or possibly the upper Cretaceous.

**Fig. 1.**
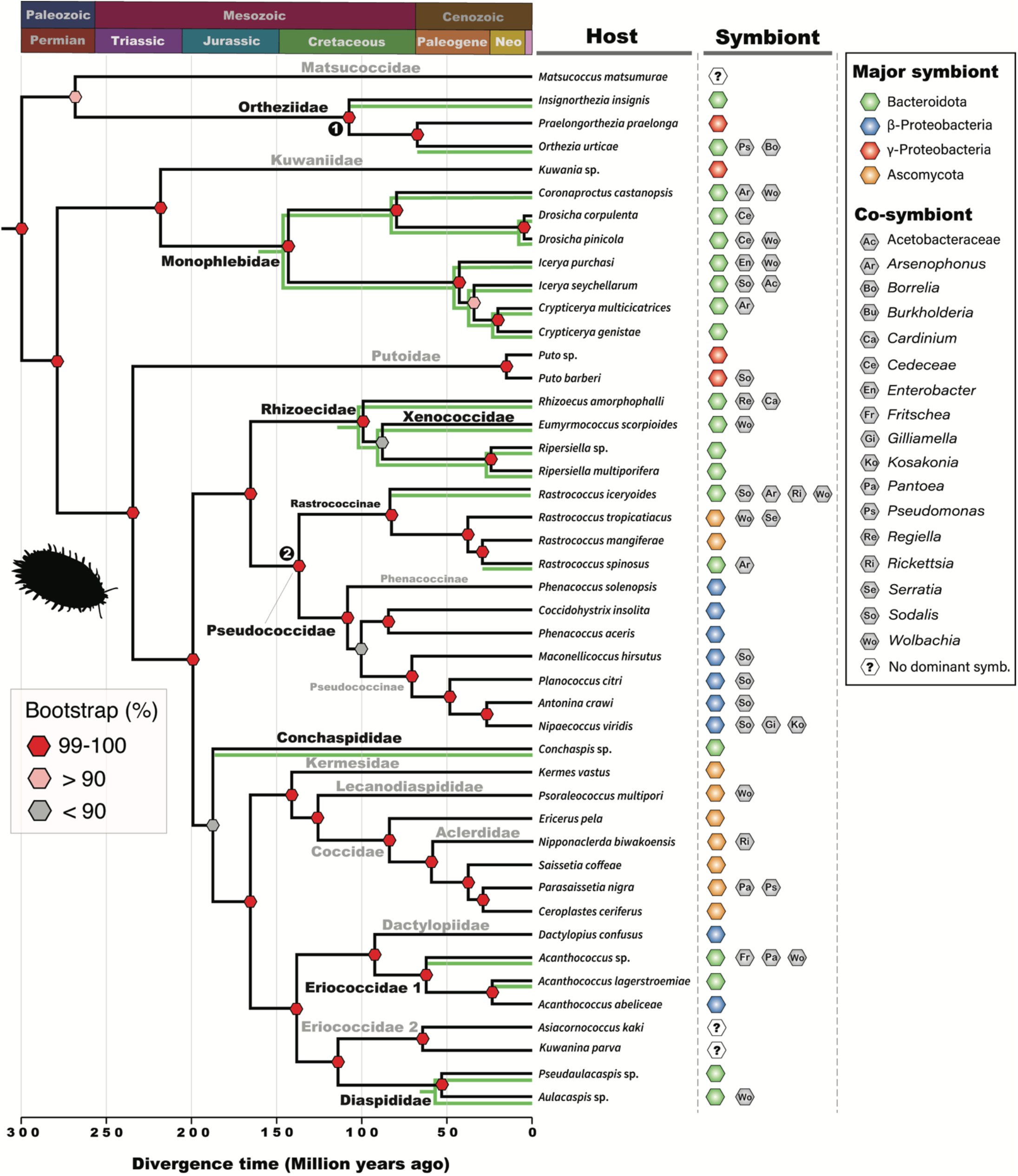
Time-estimated phylogenetic tree of scale insect hosts and their microbial symbionts. The divergence time was estimated by MCMCtree under the independent rate model with two fossil calibrations, denoted by numbered black circles. Ultrafast Bootstraps estimated by IQ-TREE are shown for each node with three different colors (red: 99-100%; pink: > 90%; gray: < 90%). Detected microbial symbionts are represented by specific symbols (see the legend on the right-hand side) next to the corresponding host species labels. The branches of host species with Bacteroidota symbionts are highlighted with shaded green lines.

### Scale insects show extreme symbiont diversity with multiple lineages of Bacteroidota symbionts

Diverse bacterial and fungal symbionts were detected from scale insects (Fig. 1). Most metagenomes (49%) included one major symbiont with a high sequence coverage and 1–4 additional co-symbionts. The remaining metagenomes contained a single symbiont (44%) or lacked a dominant symbiont (7%). Most symbionts were bacteria from the phyla Bacteroidota and Pseudomonadota (Supplementary Fig. 3). Some symbionts were fungi that mostly fell within the clade of *Ophiocordyceps* in Ascomycota (Supplementary Fig. 4). Our phylogenetic analyses support at least 10 independent origins of Bacteroidota becoming long-term endosymbionts of scale insects (Figs 1,2). All these Bacteroidota were clustered in a distinct clade with other insect symbionts within the Flavobacteriales clade, which was sister to the Weeksellaceae (Supplementary Fig. 3). In the two phylogenomic analyses that are largely congruent (Supplementary Fig. 5), most Bacteroidota symbionts of scale insects were nested within a single clade including *Skilesia*, except for *Candidatus* Brownia and Bacteroidota RAIC in a separate clade with the other insect symbionts. In the co-phylogenetic analysis, overall topologies were not mirrored between symbionts and hosts (Fig. 2; Supplementary Fig. 6). Only Bacteroidota symbionts of Diaspididae, Monophlebidae, and Rhizoecidae showed congruent topologies with their hosts. Nevertheless, due to limited sampling of free-living Bacteroidota related to the insect symbiont clade, further phylogenomics work will be needed in the future to fully resolve the exact number of independent origins of insect symbionts.

**Fig. 2.**
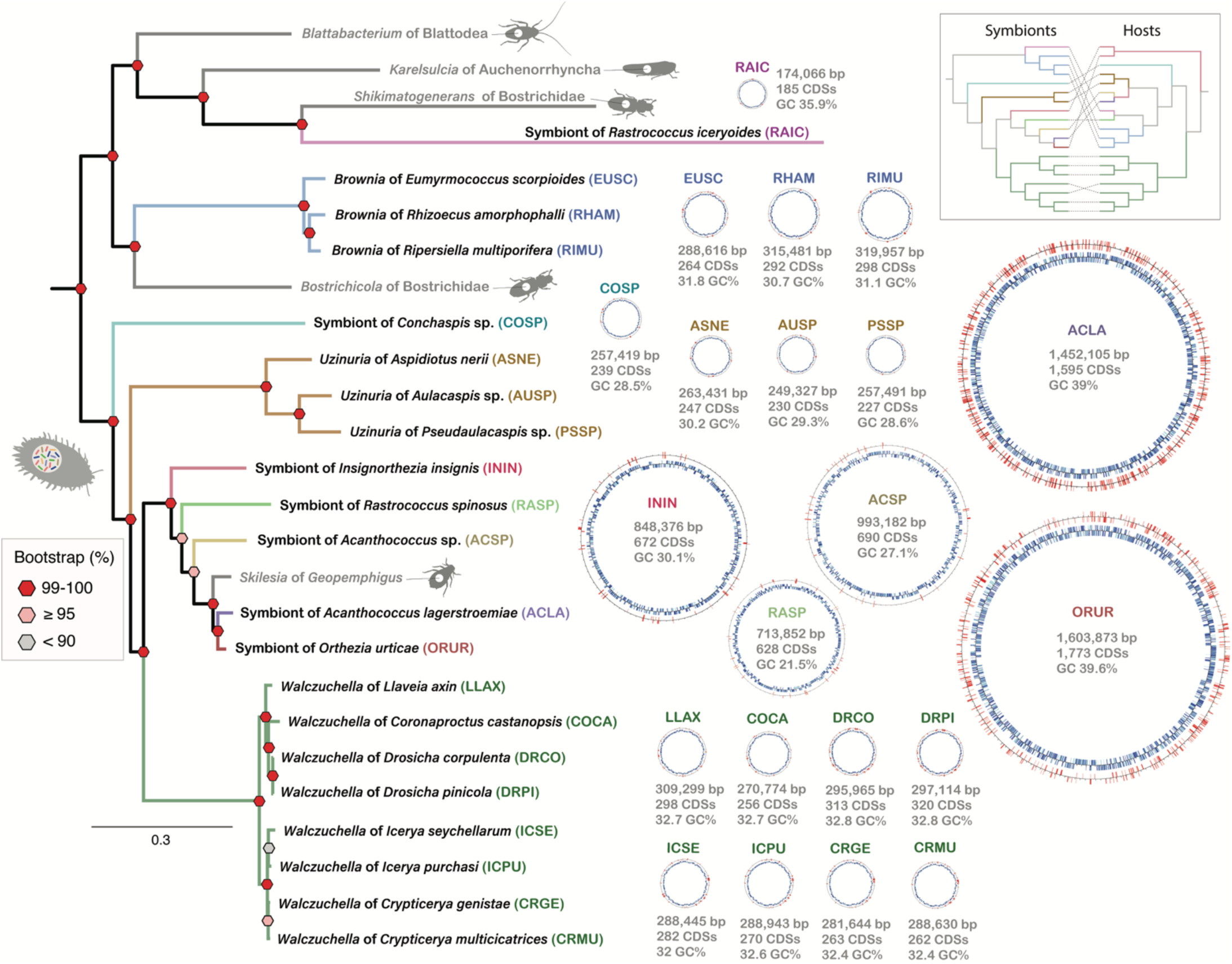
Phylogenetic relationships of Bacteroidota symbionts from scale insects with their genome features. The maximum likelihood tree was inferred with IQ-TREE based on 118 BUSCO genes under the substitution model JTTDCMut+F+R5. Ultrafast bootstraps are displayed for each node with three different colors (red: 99-100%; pink: ≥ 95%; gray: < 90%). Genome maps of 21 Bacteroidota symbionts from scale insects are placed alongside the phylogenetic tree. On genome maps, intact genes (blue) and pseudogenes (red) are represented on the inner and outer circles, respectively. Co-phylogeny of symbionts and hosts is shown on the top right.

### Multiple independent origins of the long-term Bacteroidota symbionts are reflected both in their genomes and localization

We analyzed a total of 21 genomes of Bacteroidota symbionts from scale insects, most of which were closed as complete circular-mapping genomes (Fig. 2, 3; Supplementary Table 2). The genome size of the Bacteroidota symbionts varied greatly (174–1,604 kb). The number of protein-coding genes (CDSs) of the scale insect symbionts correlated with their genome sizes, from 170 to 1,773, and coding densities from 70% to 95%. The smaller symbiont genomes (170–340 kbp) showed a higher coding density of over 79%. One exception was Bacteroidota RASP with an intermediate genome size (714 kb) but a high coding density of 88%. The genomes have GC content ranging from 21.5% to 39.6%, which surprisingly does not correlate well with their genome sizes (Fig. 2). CDSs predicted as putative pseudogenes ranged between 2–27%, with the genomes of *Walczuchella* (10– 27%), Bacteroidota ORUR (26%), and ACLA (27%) being the most highly pseudogenized. Putative plasmids of 17 kb and 3.6 kb were found in Bacteroidota ININ and COSP. Likely due to their independent origins, Bacteroidota genomes were highly rearranged and shared only limited blocks of synteny (Supplementary Fig. 7). Bacteroidota ACLA, with a large genome, is primarily distributed in the outer regions of the host’s internal body cavity and is associated with fat body cells, as shown by FISH imaging (Fig. 4A). Bacteroidota RASP, possessing an intermediate-sized genome, is intracellularly localized within host cells (Fig. 4B). These bacteriocytes are mainly found beneath the ventral host membrane. Bacteroidota RAIC is also intracellular, forming a compact organ (Fig. 4C). This bacteriome is centrally located in the host’s abdomen, near the midgut.

**Fig. 3.**
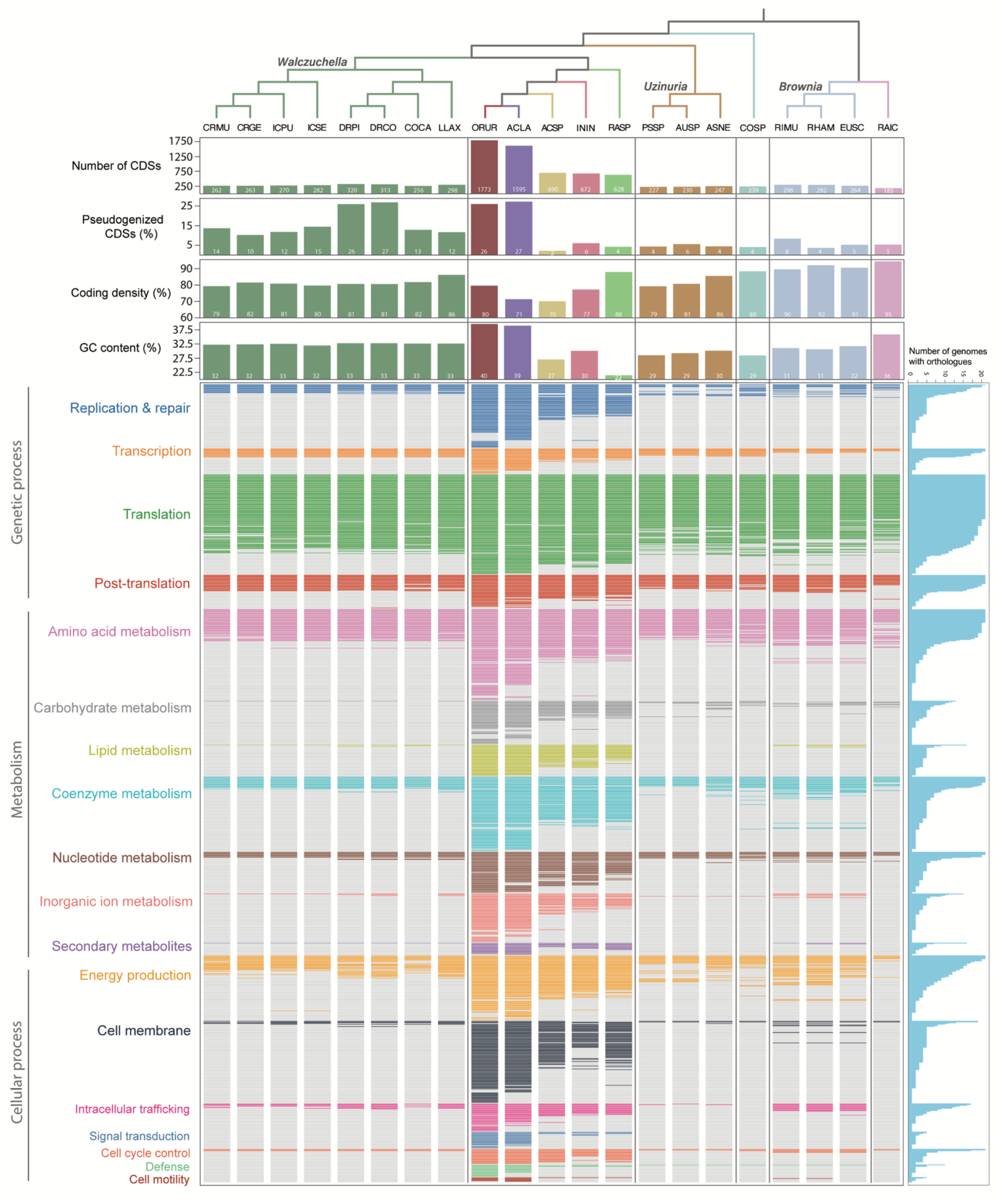
Genome properties and functional gene contents of Bacteroidota symbionts from scale insects. The bar plots indicating basic genome features are displayed above the corresponding gene content figure. Orthogroups (orthologous genes) defined with OrthoFinder were sorted into clusters of orthologous groups (COGs) by eggNOG-mapper. Colored boxes represent the presence of orthologous genes, except for pale gray color representing the absence of related genes in the genomes. The functional gene categories are listed on the left side of the main figure. The bar plot on the right indicates the number of genomes retaining corresponding orthologous genes. The phylogenetic tree represents the relationship of symbionts.

**Fig. 4.**
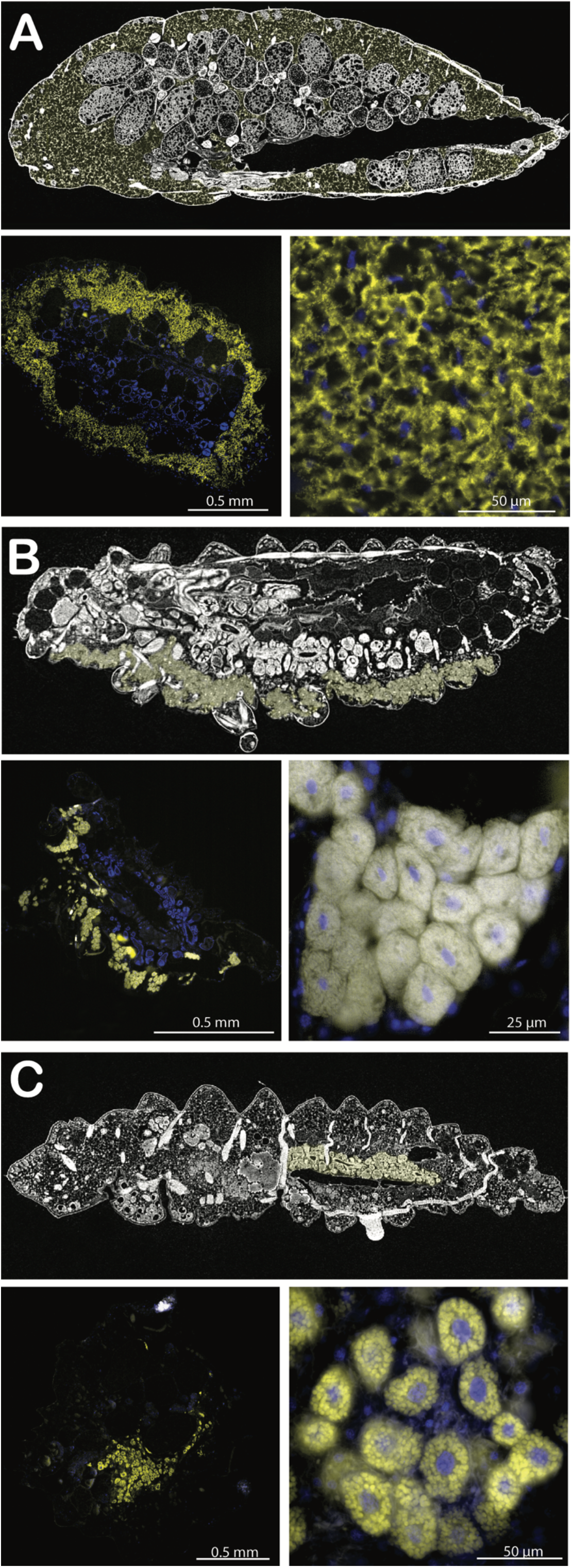
Localization of Bacteroidota in scale insects (A) *Acanthococcus largerstromae*, (B) *Rastrococcus spinosus*, and (C) *R. iceryoides*. Each panel includes body sections visualized by micro-computed tomography (µCT) (top) and fluorescence in situ hybridization (FISH) (bottom). The FISH images show whole-body sections (left) and symbiont-associated host cells (right). In the µCT images, symbiont-associated cells are highlighted in yellow. FISH images depict Bacteroidota symbionts in yellow and host cell nuclei in blue.

### Functional gene categories lost rapidly vs. more gradually

Massive gene loss has occurred in all the functional gene categories of the analyzed genomes (Fig. 3). The large symbiont genomes (with 700*–*1,600 kb genome size, hereafter, LSGs) retain 54–94% of the total orthogroups identified from all the Bacteroidota genomes. However, smaller symbiont genomes (with < 320 kb genome size, hereafter, SSGs) showed much lower retention rates of orthogroups (17–28%). Although the gene loss of Bacteroidota symbionts generally occurred in all the COGs, the protein-coding genes for translation (J), post-translational modifications (O), and AA metabolism (E) categories were proportionally increased in the SSGs compared to the LSGs (Supplementary Fig. 8).

#### Only 114 conserved core genes are retained in all the Bacteroidota symbionts of scale insects

We report 114 protein-coding genes that are retained in all the Bacteroidota symbionts of scale insects, with their *dN/dS* (ω) and predicted pseudogenes (Fig. 5A; Supplementary Table 4). Most of these core genes are related to genetic information processing (75%) and AA biosynthesis (20%). The average ω for all 114 genes was close to 0.15, excluding pseudogenes. The gene encoding 30S ribosomal subunit S20 (*rpsT*) represented the highest average ω of 0.46 among the core genes. Among symbionts, Bacteroidota RAIC showed the highest average ω of 0.28. The genes for DNA polymerase (*dnaE*) and RNA polymerase (*rpoABCD*) were intact in most symbiont genomes with 0.09–0.2 average ω values. Among translation-related genes, translation factors (*infABC, fusA, tuf*, and *tsf*) were conserved but *infC* of the two LSGs showed pseudogenization. Other conserved genes were mostly for ribosomal proteins, aminoacyl-tRNA synthetases, and post-translational processes.

**Fig. 5.**
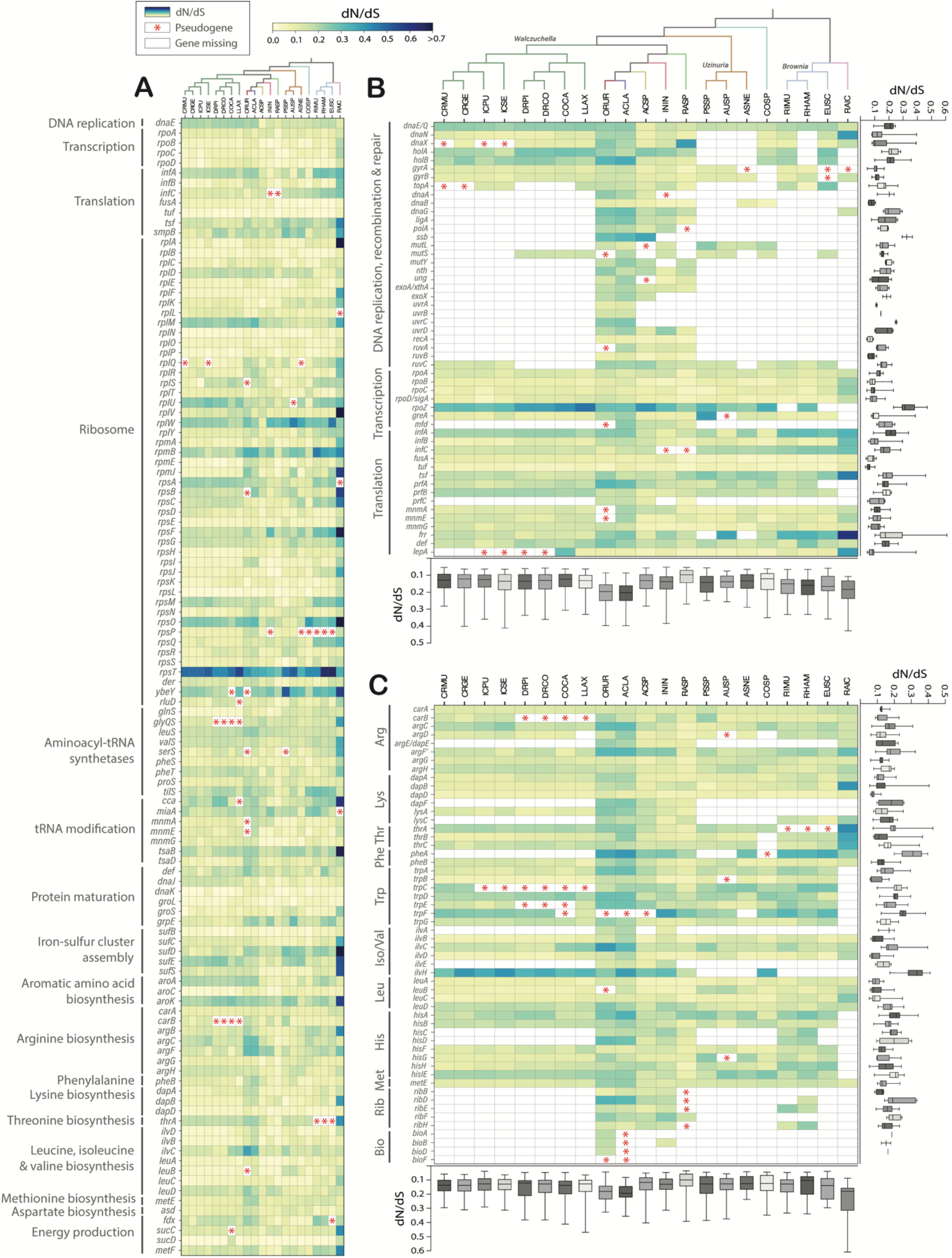
Retention pattern for selected genes of Bacteroidota symbionts from scale insects. **(A)** 114 conserved core genes of the symbionts; **(B)** Important genes for the genetic processes**; (C)** Amino acid and vitamin metabolism. Color-coding represents ratios of non-synonymous to synonymous substitutions (*dN/dS*; ω) of the retained genes. The *dN/dS* values for each gene of all the symbionts and all the retained genes of each symbiont are also shown as box plots on the right side and below the main figure, respectively. The genes flagged as pseudogenes by PseudoFinder are indicated with red asterisks (*). The missing genes are represented in white. The gene names and their related functions are listed on the left, with standard abbreviations for amino acids and B vitamins. The phylogenetic trees represent the relationships of symbionts.

#### Even core genes involved in the genetic processing and amino acid biosynthesis are lost from some genomes

We thoroughly examined the gene loss patterns in the genetic machinery and nutrition-related genes (Figs 5B-C; Supplementary Tables 5,6). The SSGs lost many key genes for DNA replication, including the replication initiation factor (*dnaA*), DNA ligase (*ligA*), DNA polymerase I (*polA*) and single-stranded DNA binding protein (*ssb*), which were absent in all of them. Although *ssb* gene is present in some LSGs, it showed the highest average ω of 0.33. In addition, genes encoding for DNA polymerase III subunits (*dnaNX* and *holAB*), DNA gyrase (*gyrAB*), helicase (*dnaB*), primase (*dnaG*) and topoisomerase (*topA*) genes are missing or pseudogenized in some SSGs. Most DNA recombination and repair genes are absent in the SSGs, and in the LSGs, many of these genes, including *mutLSY, ung, exoX*, and *uvrABC*, were also lost or pseudogenized. Among transcription-related genes, only the RNA polymerase subunit ω (*rpoZ*), transcription elongation factor (*greA*), and transcription-repair coupling factor (*mfd*) are missing or pseudogenized in some SSGs. Especially, *rpoZ* exhibited a relatively higher average ω of 0.32. The termination factor encoding gene (*prfC*) was lost from all the SSGs, as well as Bacteroidota ININ (LSG).

In addition, *prfA* and/or *prfB* were not detected in *Brownia* EUSC and Bacteroidota RAIC genomes. The SSGs showed loss and pseudogenization of at least one gene for each AA biosynthesis, with notable losses in arginine (*argE*), lysine (*dapF, dapEF*, and *lysC*), phenylalanine(*pheA*), isoleucine and valine (*ilvAEH*), and histidine (*hisCD*). Among them, *pheA* and *ilvH* showed relatively higher average ω over 0.3. Remarkably, the Bacteroidota RAIC lost all important genes for cysteine, tryptophan and histidine biosynthesis, which are likely complemented by the co-symbiont *Candidatus* Sodalis rastrocola (Supplementary Fig. 9E). Additionally, the riboflavin and biotin biosynthesis genes are missing in all the SSGs, except for a few retentions of riboflavin genes (*ribDEH*) in some *Brownia* genomes. The biotin genes were mostly missing in the LSGs as well as the SSGs. In the genetic machinery and nutritional gene categories, Bacteroidota ACLA, ORUR, and RAIC have higher average ω from 0.19 to 0.24, compared to the overall averages of 0.16 and 0.15 for these categories.

### Bacteroidota RAIC uses an alternative genetic code and lost all protein release factors

The tiny genome of Bacteroidota RAIC was inferred to use an alternative genetic code. The gene prediction for this genome with the genetic code 11 (UGA > STOP) resulted in highly fragmented protein-coding genes, in contrast to genetic code 4 (UGA > Trp) (Fig. 6A). This difference was caused by pervasive in-frame UGA codons in the open reading frames. In the genome annotation with the genetic code 4, at least one in-frame UGA codon was found in 55% of CDSs (Table 1). We also confirmed that the Bacteroidota RAIC genome has the UGA codon at the conserved positions of the *dnaE* and *valS* genes encoding tryptophan in the other symbiont genomes (Fig. 6B). The proportion of UGG encoding tryptophan was only 0.5% of the total codons across all its protein-coding sequences, which is lower than 0.7–1.0 % reported from endosymbiont genomes with the genetic code 11 (Fig. 6C). Alternatively, the UGA codon, which also encodes tryptophan, accounted for 0.3% of the codons in the Bacteroidota RAIC genome. The anticodon of tryptophan tRNA (*trnW*) is UCA in Bacteroidota RAIC, which contrasts with the other symbionts having CCA as its anticodon. The UAA and UAG represent 83% and 17% of the total stop codons in the genome of Bacteroidota RAIC (Fig. 6D). This UAA proportion was much higher than the average (64%) in the other symbionts. The Bacteroidota RAIC does not contain any genes encoding peptide release factors recognizing UAA and UAG as stop codons. Except for *Brownia* EUSC, all the other Bacteroidota symbionts retain both *prfA* and *prfB* in their genomes (Fig. 5B).

**Fig. 6.**
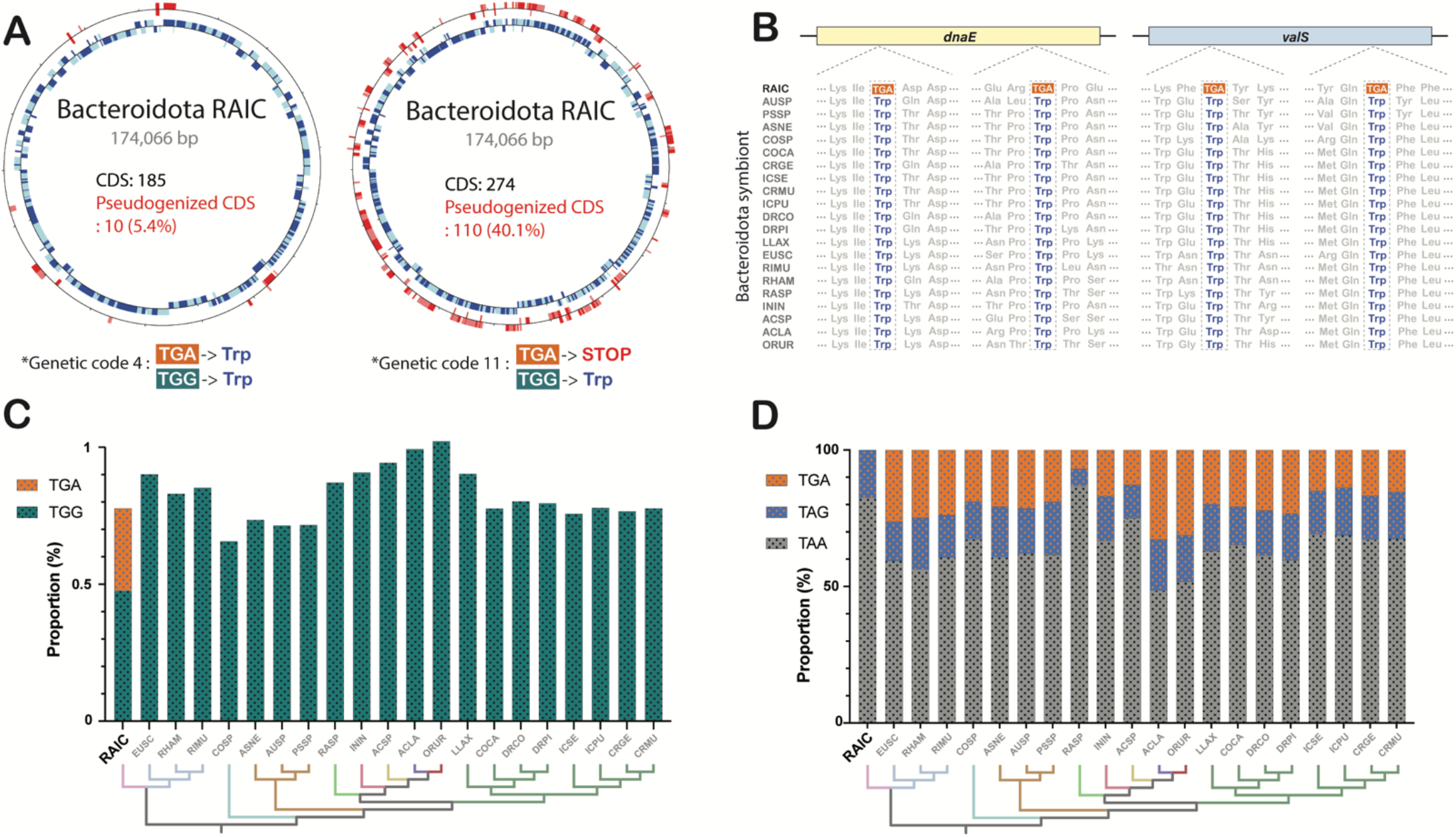
Genome features of Bacteroidota RAIC. **(A)** Comparison of genome annotations with genetic codes 4 and 11 showing different numbers of protein-coding genes (CDS) and pseudogenized CDSs. **(B)** Conserved positions encoding tryptophan in *dnaE* and *valS* genes of Bacteroidota symbionts from scale insects. Proportions of tryptophan encoded codons **(C)** and stop codons **(D)** in the symbiont CDSs. The phylogenetic trees indicate the relationships of symbionts.

## Discussion

### Bacteroidota have a long symbiotic history with insects

Our results revealed the complex evolutionary history between microbial symbionts and scale insect hosts (Fig. 1). Notably, we identified 10 different lineages of Bacteroidota that have established symbiotic relationships within eight families of scale insects. Since there were previously only five well-studied lineages of long-term Bacteroidota symbionts known from all other insect lineages, our work triples this number to 15. Based on our time-estimated scale insect phylogeny, some of the Bacteroidota symbioses have been maintained over tens to hundreds of million years such as *Brownia, Uzinuria*, and *Walczuchella* that likely entered into symbiosis with their hosts approximately 50–140 million years ago (Fig. 1). Bacteroidota COSP and RAIC may also have similarly ancient symbioses with their hosts based on their genome reduction levels (Fig. 2). On the other hand, the symbionts with larger genomes appear to have comparatively shorter relationships with their hosts (lower millions of years). The co-diversifications of *Brownia, Uzinuria*, and *Walczuchella* within individual family-level host clades suggests early interactions between different lineages of Bacteroidota and corresponding scale insects before the diversification of host lineages at the family level (Fig. 2). This contrasts with Auchenorrhyncha and cockroaches strictly co-evolving with single Bacteroidota clades *Karelsulcia* and *Blattabacterium*^15,16^. Nevertheless, given the frequency of Bacteroidota acquisitions in scale insects, it is likely that the most recent common ancestor of scale insects was associated with Bacteroidota. However, our data do not allow us to rule out other lineages, such as Pseudomonadota. Why are Bacteroidota preferred as symbionts of scale insects compared to other insect lineages? To address this fascinating question, further sampling of the free-living relatives of present-day symbionts will be needed in future studies. That no free-living or facultative Bacteroidota were identified in our sampling as loosely associated with scale insects presents a puzzling question about the ecology of the symbiotic ancestors.

### Both multiple independent acquisitions and replacements have likely shaped the current Bacteroidota symbioses with scale insects

Our findings indicate that independent acquisitions of free-living Bacteroidota lineages occurred in several scale insect sub-clades during their diversification. While some symbionts persist to the present day, others have been lost and replaced by different bacteria and fungi^27,28^. The previous hypothesis suggested host-switching of Bacteroidota symbionts from deep-branching (archeococcoids) to recently diverged (neococcoids) lineages of scale insects, primarily based on the phylogenetic similarity of the symbionts^28^. However, we consider it highly unlikely that Bacteroidota symbionts were transferred between phylogenetically distant host lineages lacking a recent common ancestor. We propose an alternative hypothesis that scale insects independently acquired specific lineages of free-living Bacteroidota, which are phylogenetically related and inclined to associate with insects. The variable genome status of scale insect Bacteroidota symbionts further suggests independent symbiotic associations rather than horizontal transmissions of symbionts among unrelated hosts. The recurrent recruitment of new microbial symbionts, rather than sustained symbiosis with a specific Bacteroidota lineage, could result from the dietary shifts of scale insects between plant phloem sap and cell contents during their evolution^76,77^. The nutrient imbalance of phloem-feeding necessitates nutritional supplementation through microbial symbiosis, in contrast to the balanced diet provided by parenchymal cells. Therefore, the frequent transitions in feeding habits are likely to lead to repeated losses and gains of microbial symbionts, ultimately contributing to the presence of multiple lineages of Bacteroidota in present-day scale insect species.

### Only genetic processing and amino acid metabolism genes are core for the Bacteroidota symbiotic with scale insects

Out of the 114 genes present in all the analyzed genomes, 93 genes have always remained intact although some are undergoing relaxed purifying selection (Fig. 5A). These genes are mainly related to genetic information processing and AA biosynthesis, highlighting the significance of these functions for the symbiont maintenance and nutrient provisioning to the insect host, as observed in other sap-feeding insects (Supplementary Fig. 8; Supplementary Fig. 10). The critical proteins involved in key genetic processes, such as DNA polymerase III subunit α, RNA polymerase subunit α and β, and σ factor, and various ribosomal proteins, remain highly conserved across Bacteroidota symbionts even from various insect hosts. The aminoacyl-tRNA synthetase genes for leucine, valine, and lysine, and tRNA modification genes also show high conservation across the studied genomes. Additionally, genes associated with post-translational processes, including SsrA-binding protein, peptide deformylase, and chaperones, are well-maintained in Bacteroidota symbionts harbored by insects. Conversely, no completely conserved genes for AA biosynthesis were identified among the insect symbiont genomes, reflecting their different gene contributions, co-symbioses with diverse partners, and nutritional requirements of their respective hosts (Supplementary Fig. 10).

### Even highly essential genes such as protein release factors were lost from some of the Bacteroidota symbionts

The multiple lineages of Bacteroidota symbionts from scale insects have experienced parallel gene losses for most functional genes, approximately representing their age (Fig. 3). This phenomenon may be attributed to the gradual elimination of redundant genes under relaxed purifying selection (Fig. 5) during long-term endosymbiosis with a certain host lineage^2,6^. In the genetic processing category, the most notable loss pertained to DNA recombination and repair genes, along with genes involved in DNA replication (Fig. 5B). The tiny symbiont genomes commonly lack certain DNA polymerase subunits, and recombination and repair genes^2^, which may contribute to increased mutation accumulation and genome instability^78,79^. While transcriptional and translational genes were generally well-preserved, certain symbiont genomes exhibited degradation in essential genes encoding translation initiation factors (IF) and peptide release factors (RF) (Fig. 5B). Notably, pseudogenization and loss of IF1 and IF2 genes occurred in some reduced or intermediate-sized genomes, affecting the proper interactions of the ribosome with mRNA^80^. Among peptide release factors, the absence of RF3 gene in most genomes probably hinders the dissociation of RF1 and RF2 from the ribosome post-peptide chain termination, influencing overall protein quantity and quality^81,82^. Although genes of RF1 and RF2, which are crucial in translation termination^83^, were relatively well preserved, their losses were still noted in two reduced genomes.

### One Bacteroidota symbiont uses an alternative genetic code

One symbiont with the most degraded genome was found to use the genetic code 4, decoding UGA to tryptophan. While this alternative genetic code is common in Mycoplasmatales, Entomoplasmatales, and mitochondria^84,85^, its occurrence is rare in animal endosymbionts. To date, only six endosymbionts, including this first case in Bacteroidota, have been found to use the genetic code 4 (Table 1). A unique feature of Bacteroidota RAIC is the absence of both *prfA* and *prfB* compared to other symbiont genomes, which at least retain intact *prfA*. This gene encodes RF1, which recognizes TAG and TAA stop codons. Hence, the mechanism by which this symbiont recognizes stop codons in translation remains uncertain. RF1 could be potentially provided by another bacterial symbiont coexisting in the same host or via its mitochondrial or cytoplasmic homologs. The loss of *prfA* may result from accidental gene loss or selective pressure induced by the TAA-biased stop codon composition in the symbiont’s coding genes (Fig. 6D). This result suggests genome ‘adaptation’ of this Bacteroidota symbiont that allowed it to avoid further gene fragmentation through severe AT mutation bias that increasingly leads to early stop codons.

### What does the frequency and evolutionary history of Bacteroidota becoming long-term endosymbionts imply for the evolution of symbiosis and the origin of organelles?

Our results demonstrate the flexibility of Bacteroidota to repeatedly transition into insect endosymbionts, as well as the preference of scale insects for symbionts from this particular bacterial clade, via a combination of sequencing and imaging data. We highlight the parallel and independent symbiont genome reduction processes together with the changes in the cell biology and localization (with the endpoint being housed in a highly specialized host organ near the midgut). The frequency with which the functionally almost identical scale insect symbioses originate and replace each other is particularly striking. Due to the rich taxon sampling of scale insects and the younger age of their symbioses (< 300 Mya) when compared to mitochondria and plastids, much more fine-grained genomics and phylogenomics analyses are feasible in this system than for organelles. The data resulting from our analyses support hypotheses such as the shopping bag model^86^, which proposes that organelles originated through successive transient endosymbioses involving multiple bacterial partners, before a stable and permanent integration. However, we highlight that even recurrently replaced symbionts often originate from phylogenetically closely related bacteria (such as one clade of Bacteroidota here). After convergently evolving for hundreds of millions of years, the symbiont genomes and cell biology are seemingly similar. The phylogenetic signal is blurred by their extreme reduction and limited by the availability of free-living relatives. Inferring the exact number of independent origins and replacements is thus more likely to lead to underestimates rather than overestimates, which should be a cautionary tale for future refinements of the shopping bag-related hypotheses.

## Acknowledgments

JYC was supported by the National Research Foundation of Korea (NRF) grant funded by the Korean government (MSIT) (2021R1A6A3A03038909) and the JSPS KAKENHI grant (20939772). FH was supported by the JSPS KAKENHI grant (23K14256) and the HFSP Early Career Grant (RGEC29/2024). We thank the Scientific Computing (SCDA), Imaging (IMG), and Sequencing (SQC) sections of the Okinawa Institute of Science and Technology for their great support.

## Data availability

The genome assemblies of Bacteroidota and *Sodalis* symbionts and Illumina raw reads are available under NCBI BioProject PRJNA1129605 and PRJNA1196661.

---

**Supplementary Figure 1.**
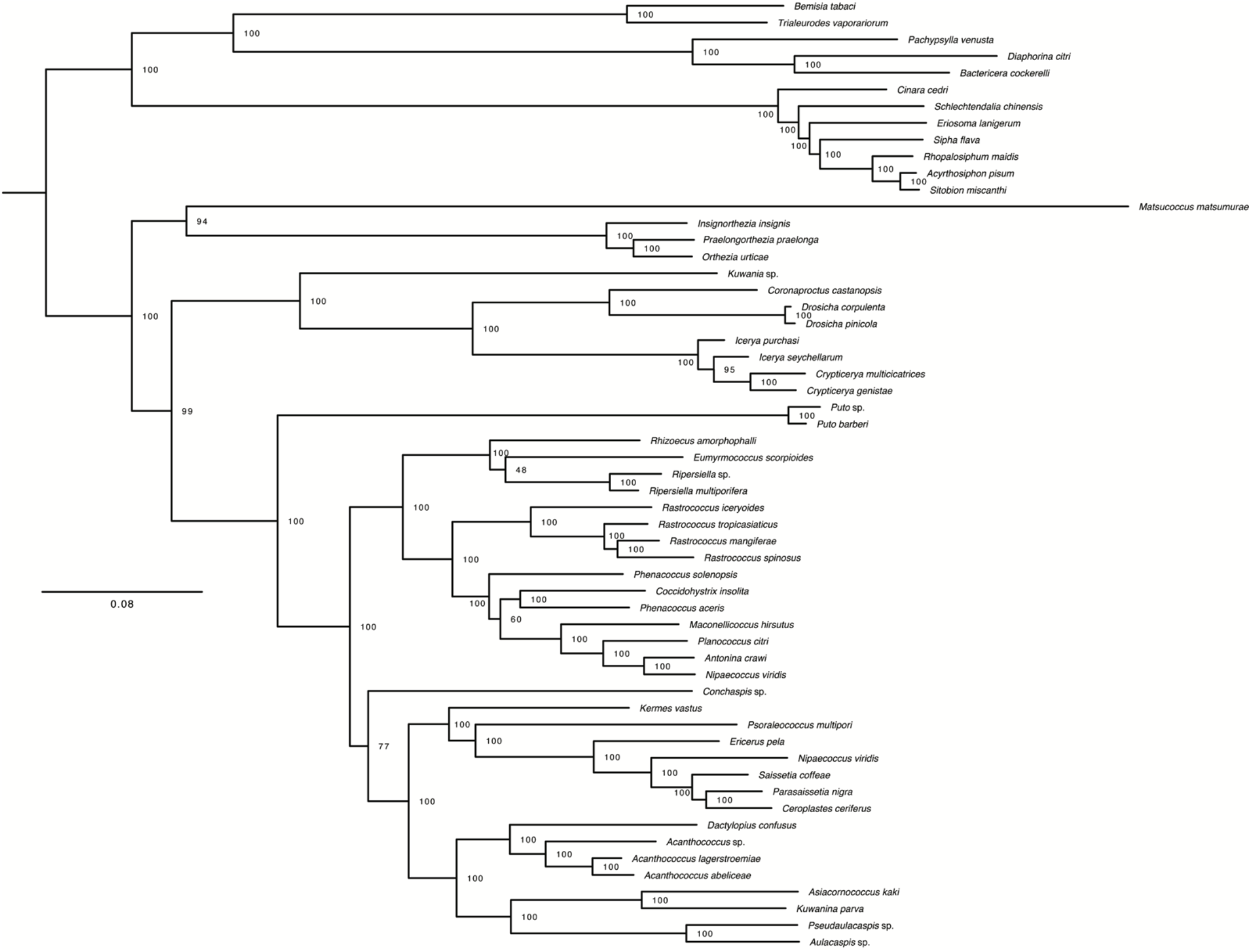
Maximum likelihood analysis of scale insect hosts inferred by IQ-TREE from 359 BUSCO genes. Numbers at each node indicate ultrafast bootstrap values.

**Supplementary Figure 2.**
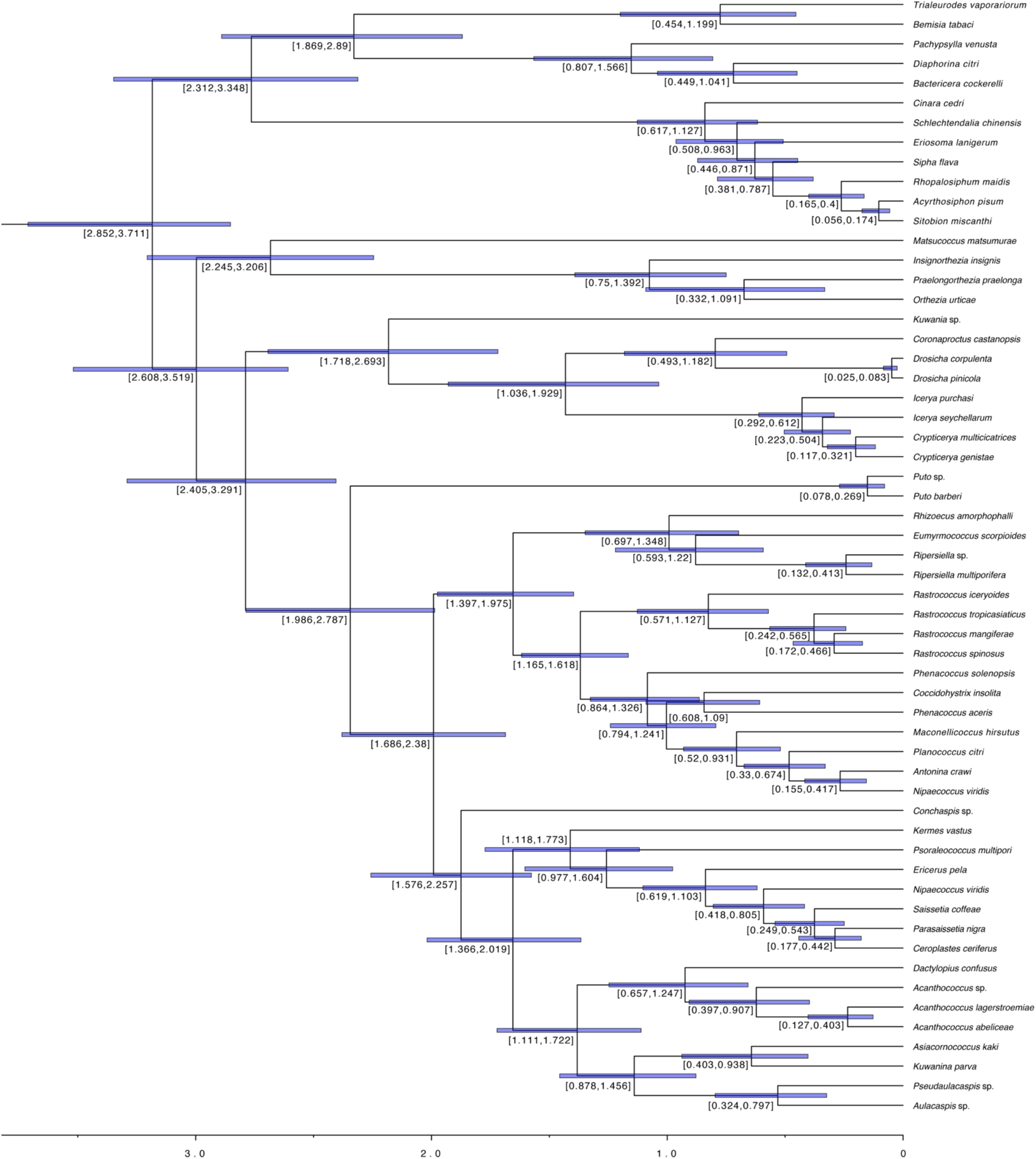
Divergence time estimates of scale insect hosts inferred by MCMCtree under the independent rate model. Numbers at each node indicate the estimated range of node age, with 95% confidence intervals displayed by blue bars. Time scale is in units of 100 million years.

**Supplementary Figure 3.**
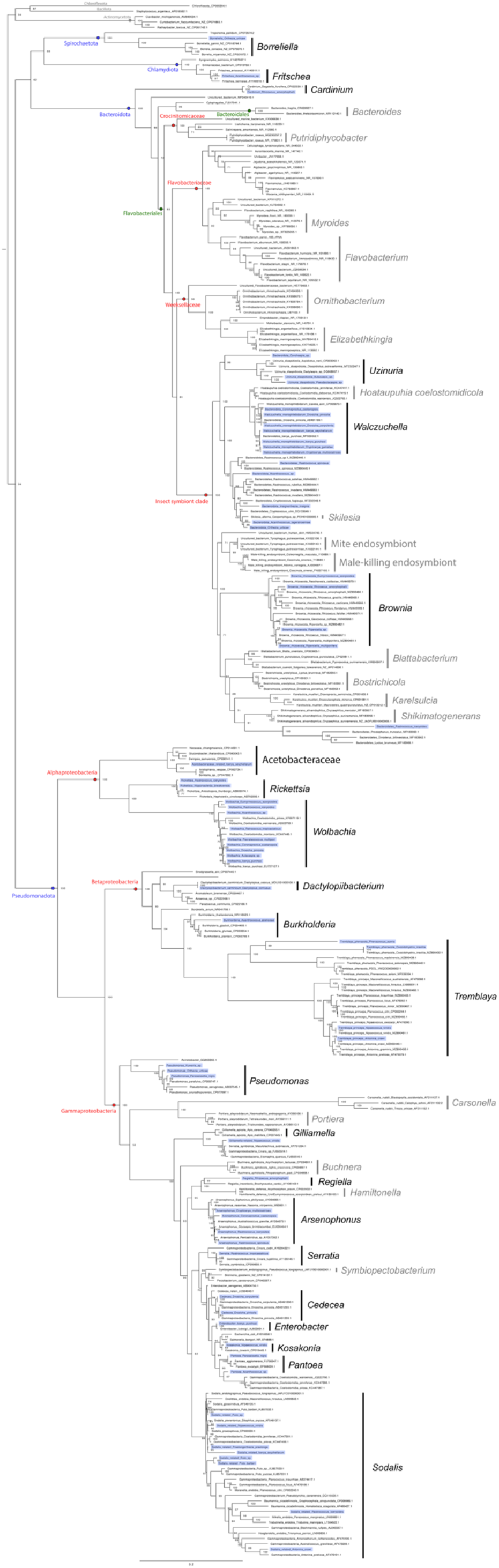
Maximum likelihood analysis of bacterial symbionts of scale insects inferred by IQ-TREE from 16S rRNA and 23S rRNA gene sequences. Numbers at each node indicate ultrafast bootstrap values. Symbionts from metagenomes used in this study are highlighted in blue.

**Supplementary Figure 4.**
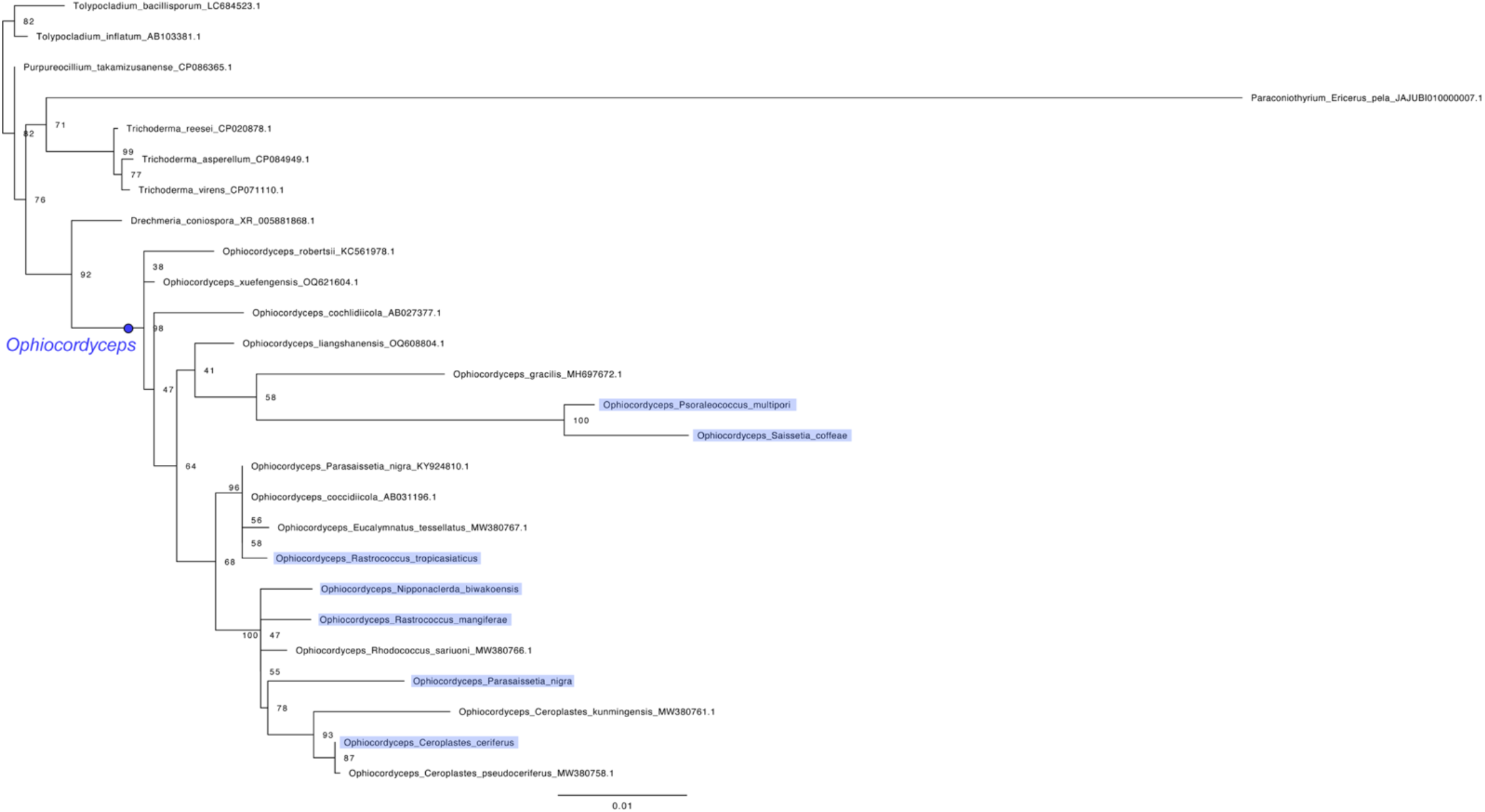
Maximum likelihood analysis of fungal symbionts of scale insects inferred by IQ-TREE from 18S rRNA and 28S rRNA gene sequences. Numbers at each node indicate ultrafast bootstrap values. Symbionts from metagenomes used in this study are highlighted in blue.

**Supplementary Figure 5.**
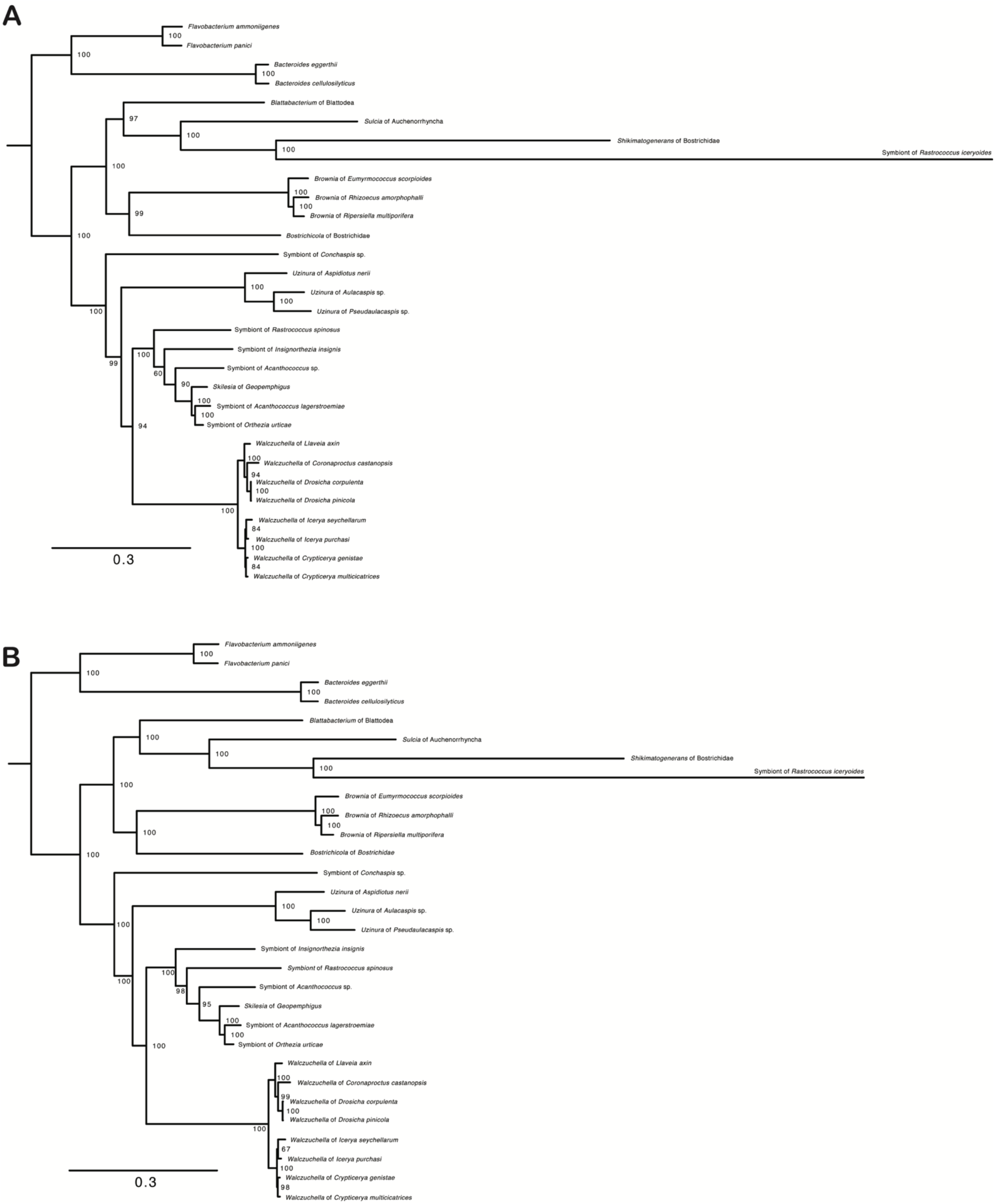
Maximum likelihood analyses of Bacteroidota symbionts from scale insects. The trees were inferred by IQ-TREE from single-copy orthologous genes sampled with OrthoFinder (65 genes) **(A)** and BUSCO (118 genes) **(B)** under the substitution models, LG+F+R5 and JTTDCMut+F+R5, respectively. Numbers at each node indicate ultrafast bootstrap values.

**Supplementary Figure 6.**
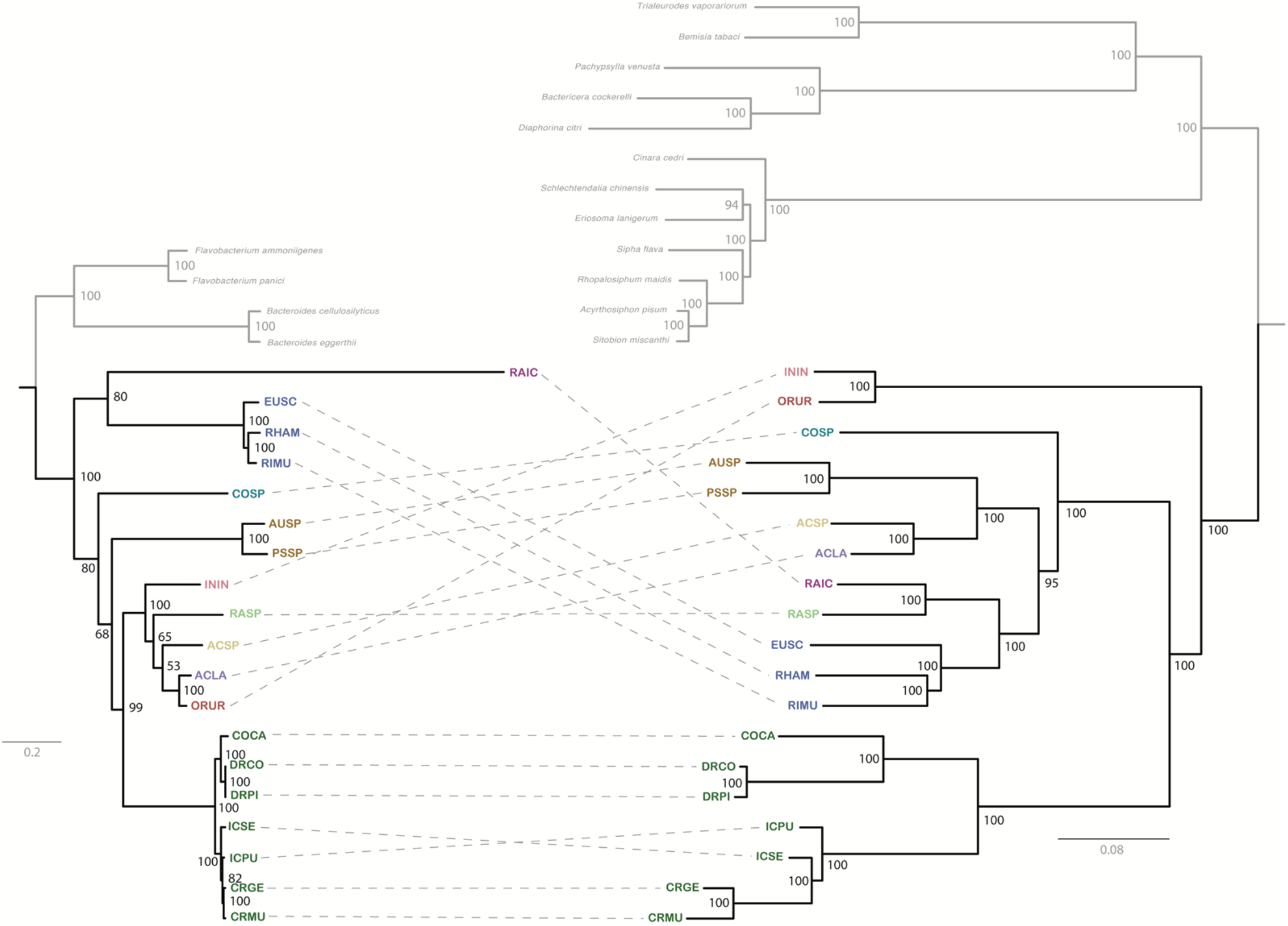
Co-phylogeny of Bacteroidota symbionts (left) and their scale insect hosts (right). The symbiont tree was constructed using 84 single-copy orthologous genes identified by OrthoFinder. The host tree was produced based on 1362 BUSCO genes. Both trees were inferred with IQ-TREE under the substitution models JTTDCMut+F+I+G4 for symbionts and LG+F+R5 for hosts. Numbers at each node indicate ultrafast bootstrap values. Symbionts are linked with dashed lines to their respective host insects. The outgroup taxa are shown in gray.

**Supplementary Figure 7.**
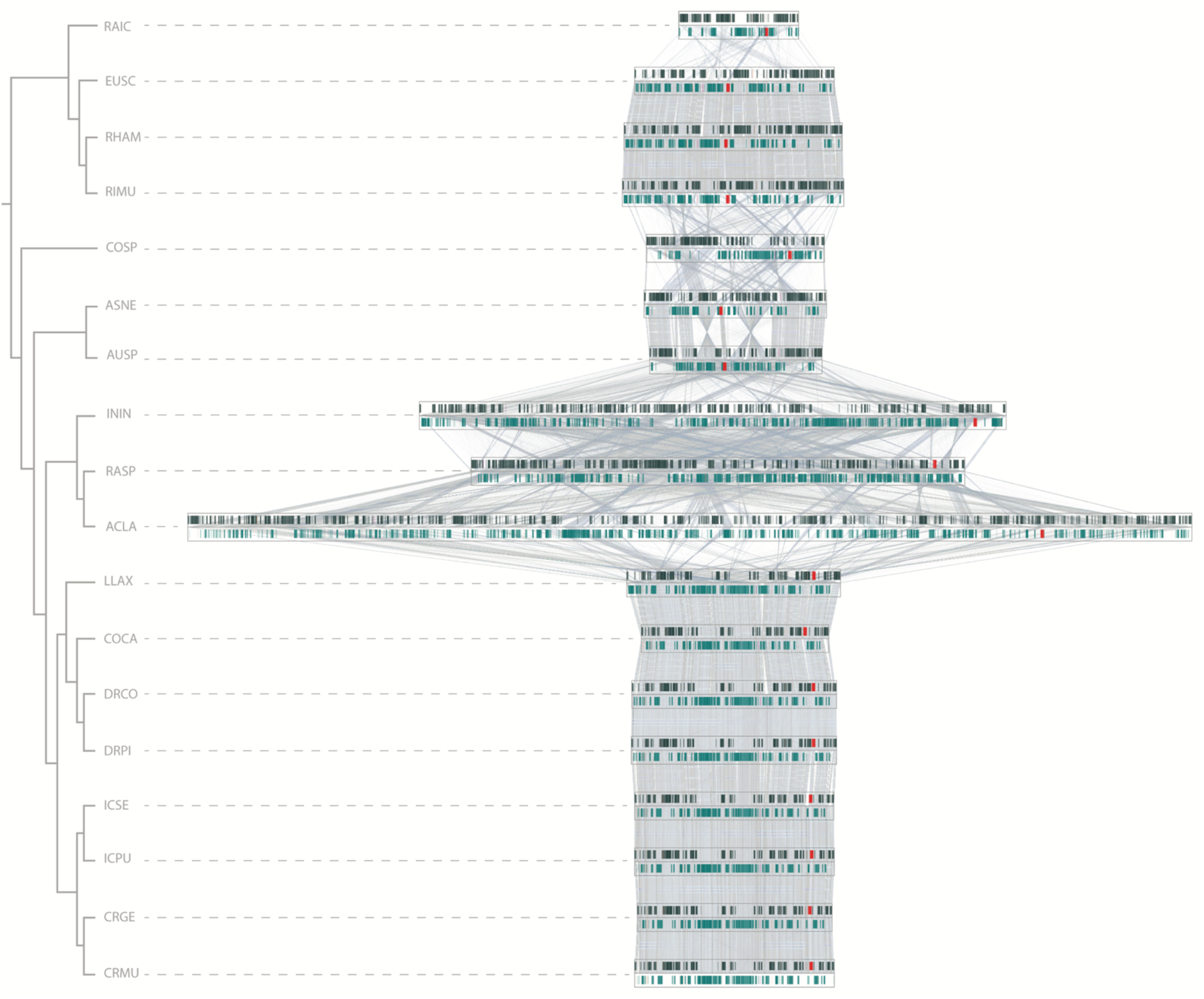
Linear genome alignments of Bacteroidota with lines linking homologous genes. While the genes of the same species are largely collinear, significant genomic rearrangements were estimated among different lineages. The genes are color-coded as follows: CDSs on the forward strand are depicted in dark grey, those on the reverse strand in cyan, rRNA in red, tRNA in orange-red, and miscellaneous RNAs in magenta.

**Supplementary Figure 8.**
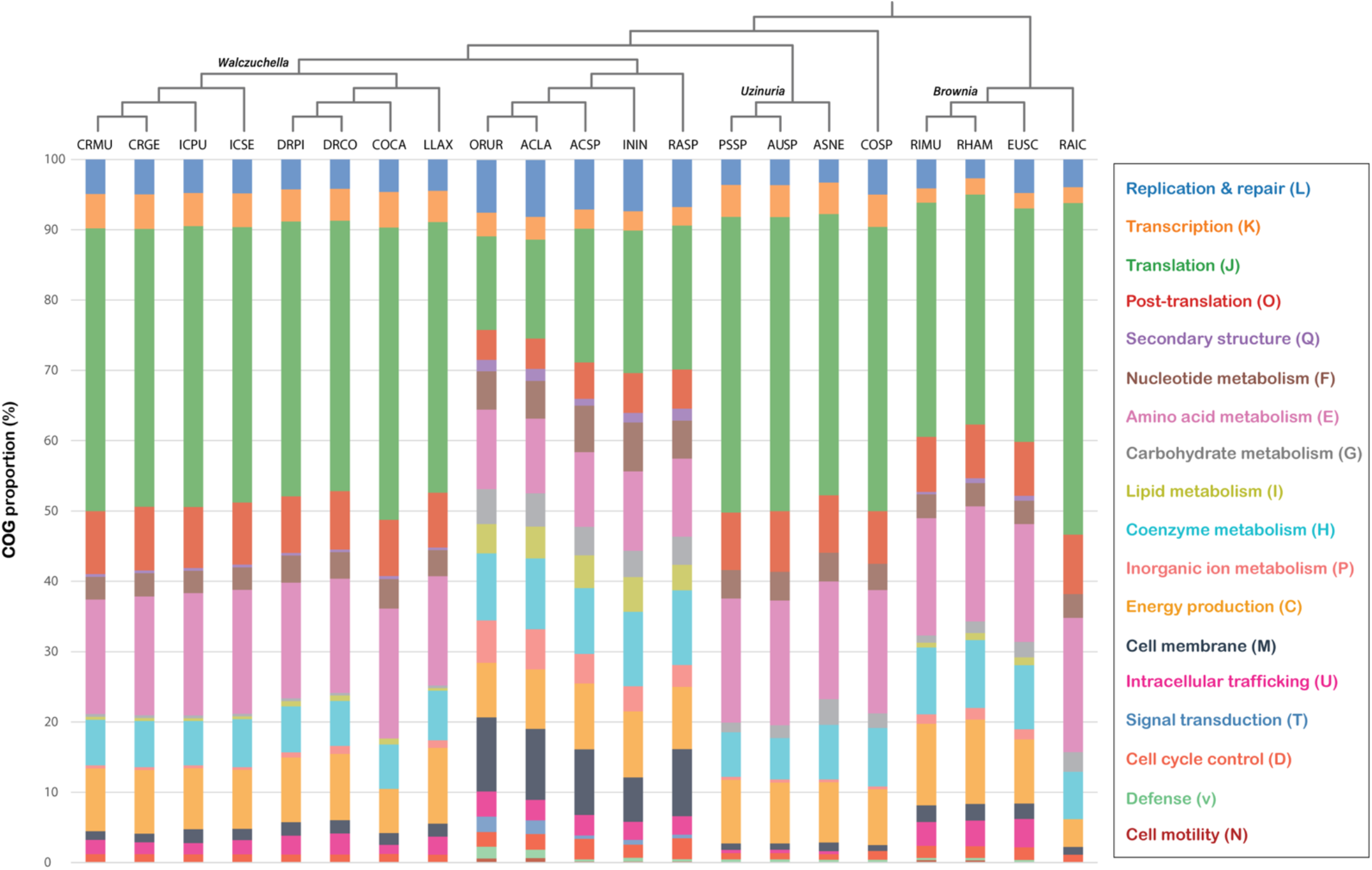
Comparison of the clusters of orthologous genes (COG) of Bacteroidota symbionts from scale insects. The legend for color coding is shown on the right.

**Supplementary Figure 9.**
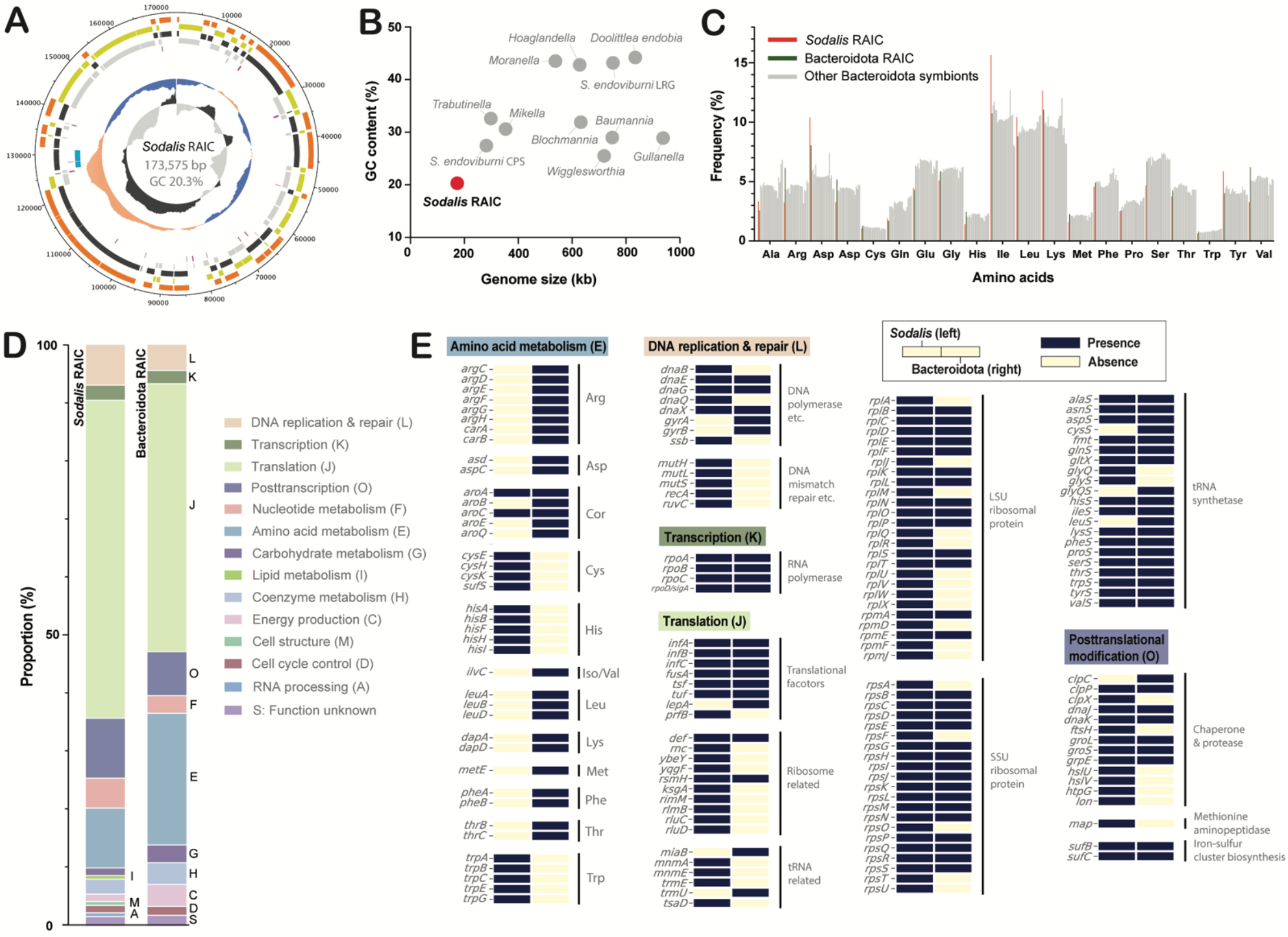
Genome properties of *Sodalis* RAIC and comparison with Bacteroidota RAIC. **(A)** Genome map of *Sodalis* RAIC. The lines from outside to inside represent: (i) CDSs on the forward strand (orange), (ii) CDSs on the reverse strand (yellow), (iii) genes on the forward strand (dark gray), (iv) genes on the reverse strand (light gray), (v) tRNA genes (purple), (vi) rRNA genes (blue), (vii) GC plot (above average with pale orange and below average with blue) and (viii) GC skew (above average with dark gray and below average with light gray). **(B)** Genome size and GC content of known *Sodalis*-like insect symbionts. **(C)** Frequency of amino acids of translated CDS sequences of symbionts. **(D)** Comparison of the cluster of orthologous genes (COG) and **(E)** gene contents for amino acid metabolism and genetic process in Bacteroidota RAIC and *Sodalis* RAIC genomes. ^*^Note: Bacteroidota RAIC is complemented by a *Sodalis*-allied co-symbiont with a tiny genome. This *Sodalis*-like symbiont was additionally found in the host of Bacteroidota RAIC (Fig. 1). This symbiont has a highly reduced genome of 173,575 bp with GC content of 20.3% (Supplementary Fig. 9A; Supplementary Table 2). Interestingly, it is the smallest genome among the known *Sodalis-*allied genomes up to date (Supplementary Fig. 9B). We name this symbiont *Candidatus* Sodalis rastrocola (hereafter, *S. rastrocola*), referring to the name of its host genus. The genome size and the number of CDSs of *S. rastrocola* are strikingly similar to the Bacteroidota RAIC genome. However, Bacteroidota RAIC retains twice as many AA biosynthesis genes as *S. rastrocola* but a much lower number of translation-related genes (Supplementary Fig. 9D). In the pathways for AA biosynthesis, the two symbionts are perfectly complementary (Supplementary Fig. 9E). For example, Bacteroidota RAIC lacks the genes for cysteine, histidine, and tryptophan, but those genes are retained in the *S. rastrocola* genome, although it lost almost all genes for the remaining AA pathways. Interestingly, all the Bacteriodota and *Sodalis*-like symbionts of scale insects show relatively low frequencies of cysteine, histidine, and tryptophan in the amino acid sequences of their protein-coding genes (Supplementary Fig. 9C). In the genetic machinery genes, *S. rastrocola* retained many genes that are missing from the Bacteroidota RAIC genome although we also notice some complementarity in the opposite direction.

**Supplementary Figure 10.**
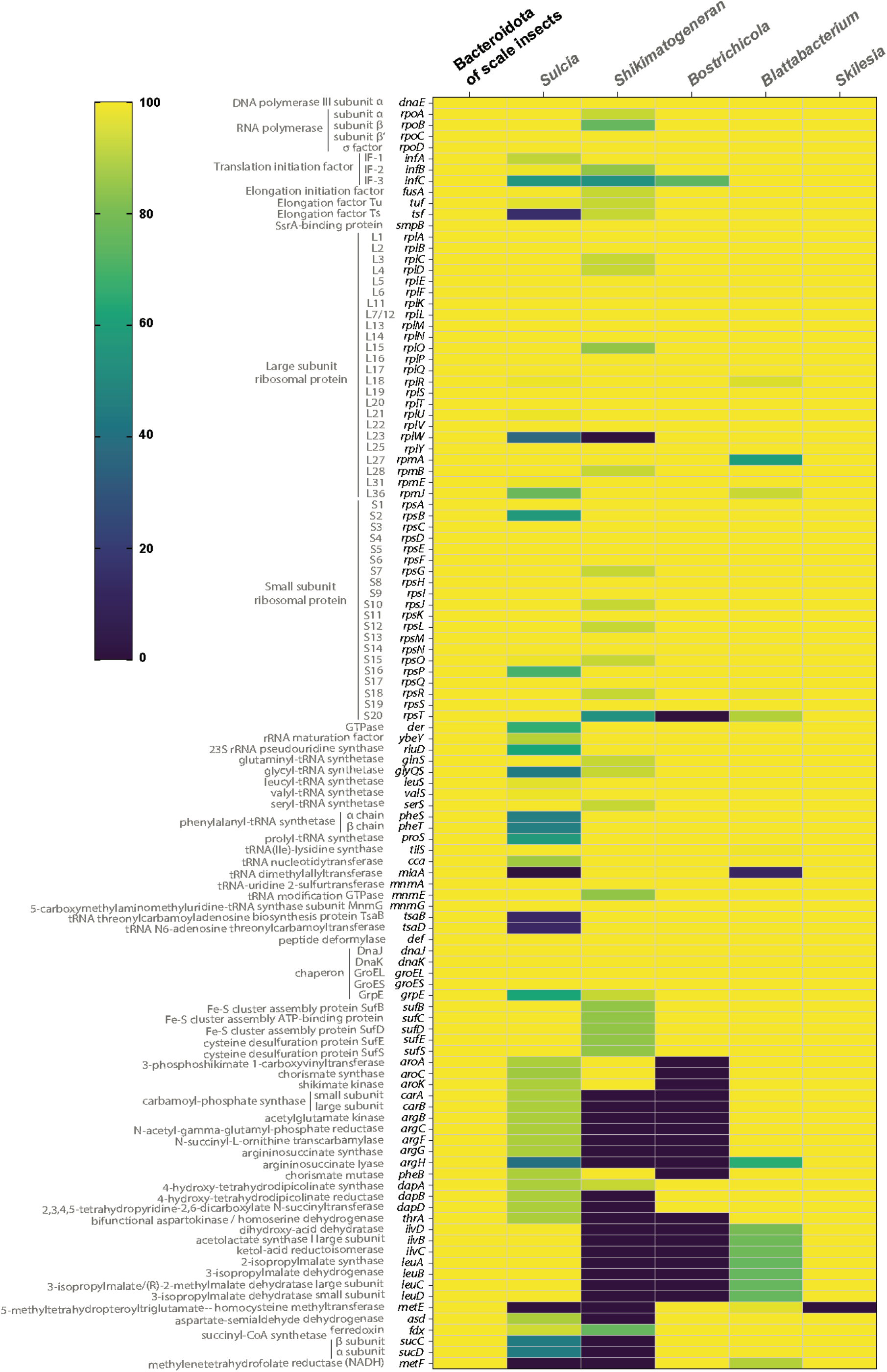
Heatmap indicating retention proportions of insect Bacteroidota symbionts for 114 conserved core genes of scale insect symbionts. Available complete genomes were used for *Blattabacterium* (#69), *Bostrichicola* (#4), *Shikimatogenerans* (#13), *Skilesia* (#1) and *Karelsulcia* (#62). The gene names, their encoded proteins, and related functions are labeled on the left. ^*^Note: Some conserved genes of scale insect symbionts are also retained in other insect symbionts (Supplementary Table 4; Supplementary Fig. 10). However, the others showed host-specific gene loss, especially in *Bostrichicola, Shikimatogenerans* and *Karelsulcia* with small genomes (142– 347 kb). Among key genes for DNA replication, transcription and translation, *dnaE* and *rpoCD* are present in all genomes of other insect symbionts. In contrast, the missing *infC* was confirmed in the three small genome symbionts above. In addition, *tsf* is lacking in most *Karelsulcia* genomes. Almost all genes for ribosome biosynthesis are retained in *Blattabacterium, Bostrichicola* and *Skilesia* genomes, whereas *Shikimatogenerans* and *Karelsulcia* lack some of the ribosomal biosynthesis genes in their genomes. In particular, *rplW* is missing in all the genomes of *Shikimatogenerans*. For the tRNA biosynthesis genes, two genes (*mnmAG*) are completely maintained in all the genomes of other insect symbionts. However, *miaA* and *tsaBD* are absent in almost all genomes of *Karelsulcia*, which also lack other tRNA biosynthesis genes. Most genomes of *Blattabacterium* also lost *miaA*. Among the genes for post-translational processes, the gene for rescuing stalled ribosomes (*smpB*) and protein maturation-related genes (*def, dnaJK* and *groLS*) are retained in all the insect symbiont genomes. The genes for iron-sulfur cluster biosynthesis (*sufBCDE*) are present in all the genomes of other insect symbionts except for some genomes of *Shikimatogenerans*. The AA biosynthesis genes showed clear missing in *Bostrichicola* and *Shikimatogenerans* genomes. *Karelsulcia* genomes well retain genes for the biosynthesis of cysteine, (iso)leucine and valine (*ilvBCD, leuABCD* and *sufS*), but some genomes lost the other AA genes. In contrast, *Blattabacterium* showed a specific loss for the (iso)leucine and valine biosynthesis genes in some genomes, as well as *argH*. The energy production-related genes are relatively conserved in *Bostrichicola, Blattabacterium* and *Skilesia* genomes, but the clear missing *sucCD* and *metF* were confirmed in the genomes of *Shikimatogenerans* and *Karelsulcia. Skilesia* genome retains almost all the conserved genes of scale insect symbionts, except for *metE*.

